# Orientation-dependent *Dxz4* contacts shape the 3D structure of the inactive X chromosome

**DOI:** 10.1101/165340

**Authors:** G. Bonora, X. Deng, H. Fang, V. Ramani, R. Qui, J. Berletch, G. N. Filippova, Z. Duan, J. Schendure, W. S. Noble, C. M. Disteche

## Abstract

The mammalian inactive X chromosome (Xi) condenses into a bipartite structure with two superdomains of frequent long-range contacts separated by a boundary or hinge region. Using in situ DNase Hi-C in mouse cells with deletions or inversions within the hinge we show that the conserved repeat locus *Dxz4* alone is sufficient to maintain the bipartite structure and that *Dxz4* orientation controls the distribution of long-range contacts on the Xi. Frequent long-range contacts between *Dxz4* and the telomeric superdomain are either lost after its deletion or shifted to the centromeric superdomain after its inversion. This massive reversal in contact distribution is consistent with the reversal of CTCF motif orientation at *Dxz4.* De-condensation of the Xi after *Dxz4* deletion is associated with partial restoration of TADs normally attenuated on the Xi. There is also an increase in chromatin accessibility and CTCF binding on the Xi after *Dxz4* deletion or inversion, but few changes in gene expression, in accordance with multiple epigenetic mechanisms ensuring X silencing. We propose that *Dxz4* represents a structural platform for frequent long-range contacts with multiple loci in a direction dictated by the orientation of a bank of CTCF motifs at *Dxz4,* which may work as a ratchet to form the distinctive bipartite structure of the condensed Xi.

---

Mammalian X chromosome inactivation (XCI) results in the silencing of one of the two X chromosomes in female somatic cells, which is initiated by expression of the lncRNA *Xist* from the inactive X chromosome (Xi). This is followed by cis-spreading of *Xist* RNA and recruitment of specific proteins that implement repressive epigenetic changes, including loss of RNA polymerase II and of active histone marks such as H3-H4 acetylation and H3 methylation at lysine 4, as well as enrichment in silencing marks such as H3 tri-methylation at lysine 27 (H3K27me3) and H2A ubiquitination at lysine 119. These initial epigenetic modifications are followed by accumulation of the histone variant macroH2A, tri-methylation of H3 at lysine 9 and of H4 at lysine 20, together with DNA methylation at CpG islands^1,2^. Drastic changes in the structure of the Xi—most visibly condensation—take place once it becomes silenced. Early microscopy studies reported a condensed chromatin body (Barr body) specifically present in female cells, and later shown to represent the Xi, often located at the nuclear periphery or adjacent to the nucleolus^3-5^.

Genome-wide chromosome conformation capture (Hi-C) studies in human and mouse cells and tissues demonstrate that chromosomes are divided into two types of compartments, A and B, associated with open and closed chromatin, respectively^6^. In contrast, allelic contact maps for the condensed human and mouse Xi show two superdomains of contacts separated by a hinge, forming a characteristic bipartite 3D structure^7-11^. Long-range contacts are frequent within each superdomain, but are not observed between superdomains, and there is little evidence of A/B compartments as compared to the Xa or the autosomes. While the genomic content of the superdomains differs between human and mouse, the hinge region is partially preserved and contains the macrosatellite repeat locus *DXZ4/Dxz4* in both species^7-11^. The *DXZ4/Dxz4* loci encode lncRNAs and bind CTCF and components of the ring-shaped cohesin complex only on the Xi, while the loci are methylated on the active X (Xa), preventing CTCF binding^12-16^. CTCF and cohesin are two of the main organizers of nuclear structure^17-20^. New studies have shown that highly dynamic chromatin loops are formed by progressive extrusion of the chromatin fiber due to the movement of extruding factors (EFs) such as cohesin rings. Extrusion proceeds until a boundary element (BE) bound to chromatin, such as CTCF, is encountered, which stalls the loop and ultimately defines topologically associated domains (TADs)^21,22^. Within a TAD, loops can be continuously formed by loading of cohesin by the SCC2/SCC4 complex, processive loop extension, and release by WAPL^23,24^. Convergent CTCF binding motifs (i.e. facing each other) at the base of a loop favor strong interactions and the inversion of CTCF sites disrupts loop formation^11,25,26^. In the case of *DXZ4/Dxz4* CTCF motifs are arranged in tandem orientation at the locus, with an estimated 10-100 copies in human^14^, and 14 copies in mouse^12^, which was confirmed using Blast alignment of a 34bp conserved CTCF binding sequence. How the CTCF motif arrangement influences long-range chromatin contacts on the Xi is unknown. In addition to *Dxz4,* the mouse hinge region originally defined using Hi-C also contains the mouse-specific minisatellite repeat *Ds-TR* whose function is unknown^7,12^. Both *Dxz4* and *Ds-TR* loci bind nucleophosmin, an essential component of the nucleolus, and could represent a large nucleolus-associated domain (NAD) that may help position the Xi near the nucleolus^7,15^.

To determine the role of each element of the hinge including *Dxz4* and *Ds-TR* in the maintenance of the 3D structure of the mouse Xi in relation to its silencing and nuclear positioning in somatic cells, we used allele-specific CRISPR/Cas9 editing to induce different sized deletions and inversions specifically targeted to the Xi. We then tested effects of these modifications on the overall 3D structure of the Xi using *in situ* DNase Hi-C^27^. High resolution allele-specific analyses were done to assess changes in the distribution of contacts and in the TAD structure along the X chromosomes. We scored these changes in relation to CTCF binding profiles obtained by ChIP-seq and to chromatin accessibility profiles obtained by ATAC-seq. We determined the effects of genomic alterations of the hinge on the size and position of the Xi in the nucleus relative to the nuclear periphery and the nucleolus. Finally, gene expression changes were measured by RNA-seq, considering genes normally silenced by XCI and genes that escape XCI.

## Results

### 1. The integrity of the superdomains on the mouse Xi depends on *Dxz4,* but not *Ds-TR*

To evaluate the role of specific elements located within the hinge that separates superdomains of long-range intrachromosomal interactions on the mouse Xi, we used allele-specific CRISPR/Cas9 editing to induce genomic alterations in Patski fibroblast cells, in which skewed XCI and frequent species-specific polymorphisms (1/93bp using validated Sanger MM10 *Mus spretus* SNPs) allowed us to distinguish the Xi from C57BL/6 (BL6) and the Xa from *Mus spretus*^7,28,29^. Xi-specific deletions or inversions were induced using pairs of small guide RNAs (sgRNAs) that flank either most of the hinge region or only some of its elements (Supplementary Table S1). We isolated two independent single-cell clones (hereafter, Del-hinge clone a and b) with a large 127kb deletion of the hinge that includes both *Dxz4* and *Ds-TR* (Supplementary Fig. S1a). We also derived an independent single-cell clone with a 44kb deletion of *Dxz4* alone (hereafter Del-Dxz4), two independent clones with a 44kb inversion of *Dxz4* (hereafter Inv-Dxz4 clone a and b), and single clones with a 37kb deletion of *Ds-TR* (hereafter Del-Ds-TR) alone, or a small 907bp inversion of 2 of 3 CTCF binding sites located at the 5’end of *Ds-TR* (hereafter Inv-5’ Ds-TR) (Supplementary Fig. S1a). Del-hinge clone a and Inv-Dxz4 clone a were used for Hi-C, ChIP-seq, ATAC-seq and microscopy analyses, whereas both clones a and b were used for RNA-seq. Sanger DNA sequencing analyses verified that the genomic alterations obtained by CRISPR/Cas9 editing were specific to the Xi allele, and Del-hinge clone a was also confirmed by FISH (Supplementary Fig. S1b, c and data not shown).

Next, in situ DNase Hi-C of the edited cell clones was performed in comparison to wild-type Patski cells using an established method (Supplementary Table S2)^7,27^. Contact maps for the Xi showed that a large deletion incorporating both *Dxz4* and *Ds-TR* (Del-hinge) as well as a deletion of *Dxz4* alone (Del-Dxz4) dramatically disrupted the bipartite structure of the Xi normally observed in wild-type cells (WT) (Fig. 1a, b). In contrast, deletion of *Ds-TR* (Del-Ds-TR) alone or alteration of its 5’end (Inv-5’ Ds-TR) did not affect the Xi bipartite structure or the contact distribution. There was no apparent change in the contact maps of the Xa or of the autosomes in any of the deleted/inverted cell clones (Supplementary Fig. S2a). The bipartite structure of the Xi was clearly abrogated by either deletion of *Dxz4* alone or by deletion of the hinge including both *Dxz4* and *Ds-TR.* We noted subtle differences in the contact maps between these two mutations, with the contact map obtained after deletion of the hinge resembling more closely that of the Xa with respect to A/B compartment structure (Fig. 1a and Supplementary Fig. S2a). Thus, we cannot exclude that *Ds-TR* or another hinge element plays a minor role in the Xi structure in the context of a *Dxz4* deletion, even though the *Ds-TR* deletion by itself did not cause any apparent change to the contact map of the Xi. Inversion of *Dxz4* was associated with persistence of the Xi bipartite structure, but caused extensive re-distribution of contacts as described below. We conclude that *Dxz4* alone is necessary for the formation of the two superdomains on the Xi.

**Fig. 1:**
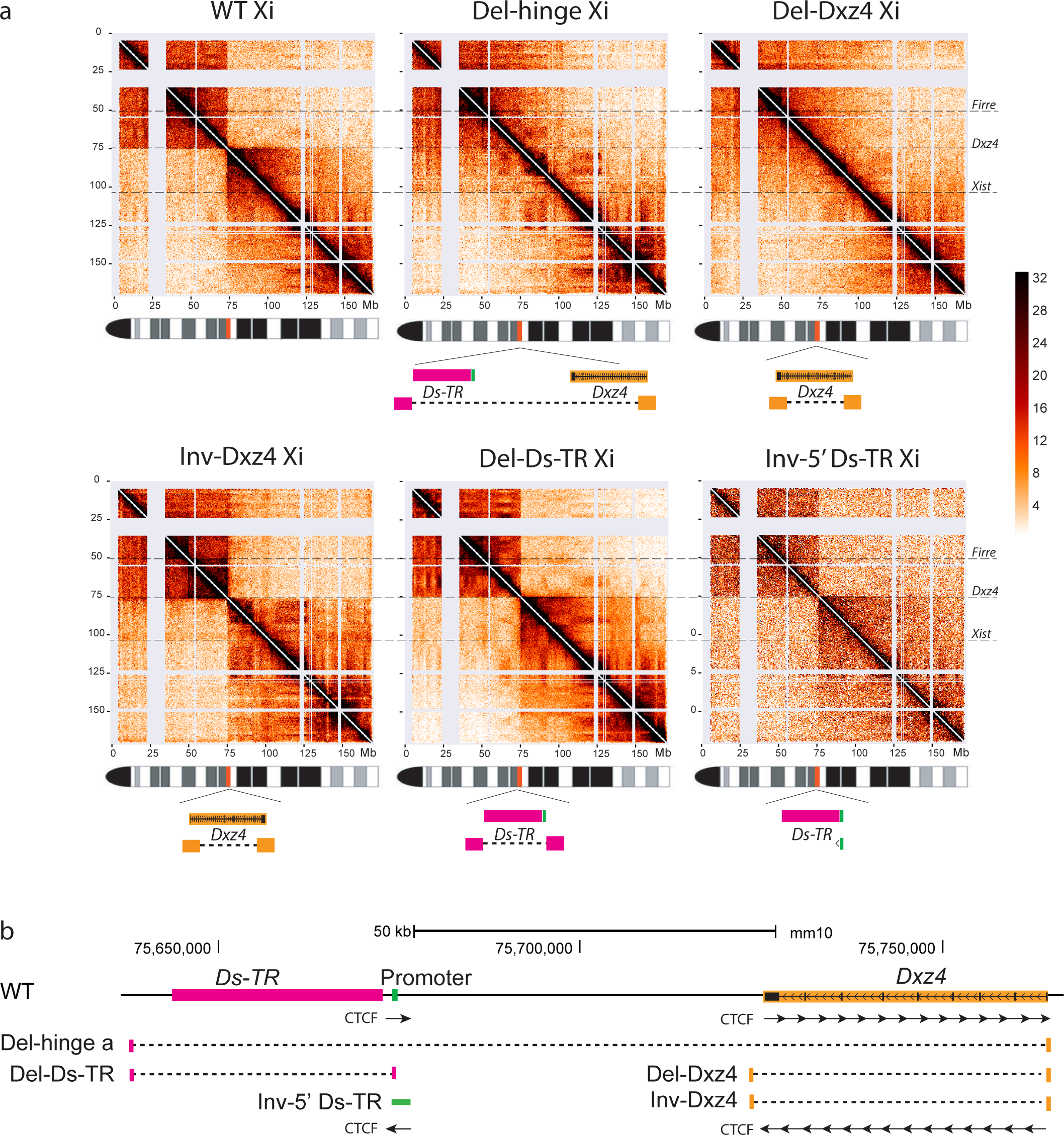
*Dxz4* alone is sufficient to maintain the bipartite structure of the Xi. **a**. Contact maps are shown at 500kb resolution for the Xi in WT, Del-hinge (Xi deletion nt75637519-75764753), Del-Dxz4 (Xi deletion nt75721096-75764754), Inv-Dxz4 (Xi inversion nt75721096-75764754), Del-Ds-TR (Xi deletion nt75637501-75674037), and Inv-5’ Ds-TR (Xi inversion nt75674046-75674952). Note that fewer reads were obtained for the Inv-5’ Ds-TR, resulting in lower resolution (Supplementary Table S2). The location of *Dxz4, Firre* and *Xist* and schematics of the allele-specific deletions/inversions are shown. The color scale shows normalized contact counts. See Supplementary Fig. S2a for contact maps of the Xa in the same clones. **b**. Relative position of the loci within the hinge region and location of the CRISPR/Cas9 induced alterations. Arrows indicate the orientation of CTCF motifs at *Dxz4*^11^ and at the 5’end of *Ds-TR.* See Fig. 5b for CTCF binding at *Dxz4* on the Xi by ChIP-seq.

### 2. *Dxz4* contacts with loci in the telomeric superdomain are disrupted by *Dxz4* deletion or inversion

To increase the total number of allelic reads for better resolution, two pooled sets of Hi-C contacts were generated: a data set representing Del-hinge and Del-Dxz4 (designated Del-hinge/Dxz4) and another representing wild-type and Del-Ds-TR(designated WT*) (Supplementary Table S2). Allelic contact maps for each individual data set are very similar to those obtained from the pooled data in terms of the presence or absence of the bipartite structure, justifying the pooling (Fig. 1a; Supplementary Fig. S2a, b). Raw and Pearson correlation-transformed contact maps of the Xi clearly show that the hinge region, which is present in WT*, almost disappears in Del-hinge/Dxz4 (Fig. 2a; Supplementary Fig. S2b). Zooming in to examine a 50Mb region around *Dxz4* clearly shows the loss of the separation between superdomains in Del-hinge/Dxz4, and the strong shift in contacts in Inv-Dxz4 with new contacts appearing between *Dxz4* and *Firre,* another macrosatellite repeat locus that also binds CTCF on the Xi^15,30^ (Fig. 2b). As expected, no hinge is seen on the Xa (Supplementary Fig. S2b).

**Fig. 2:**
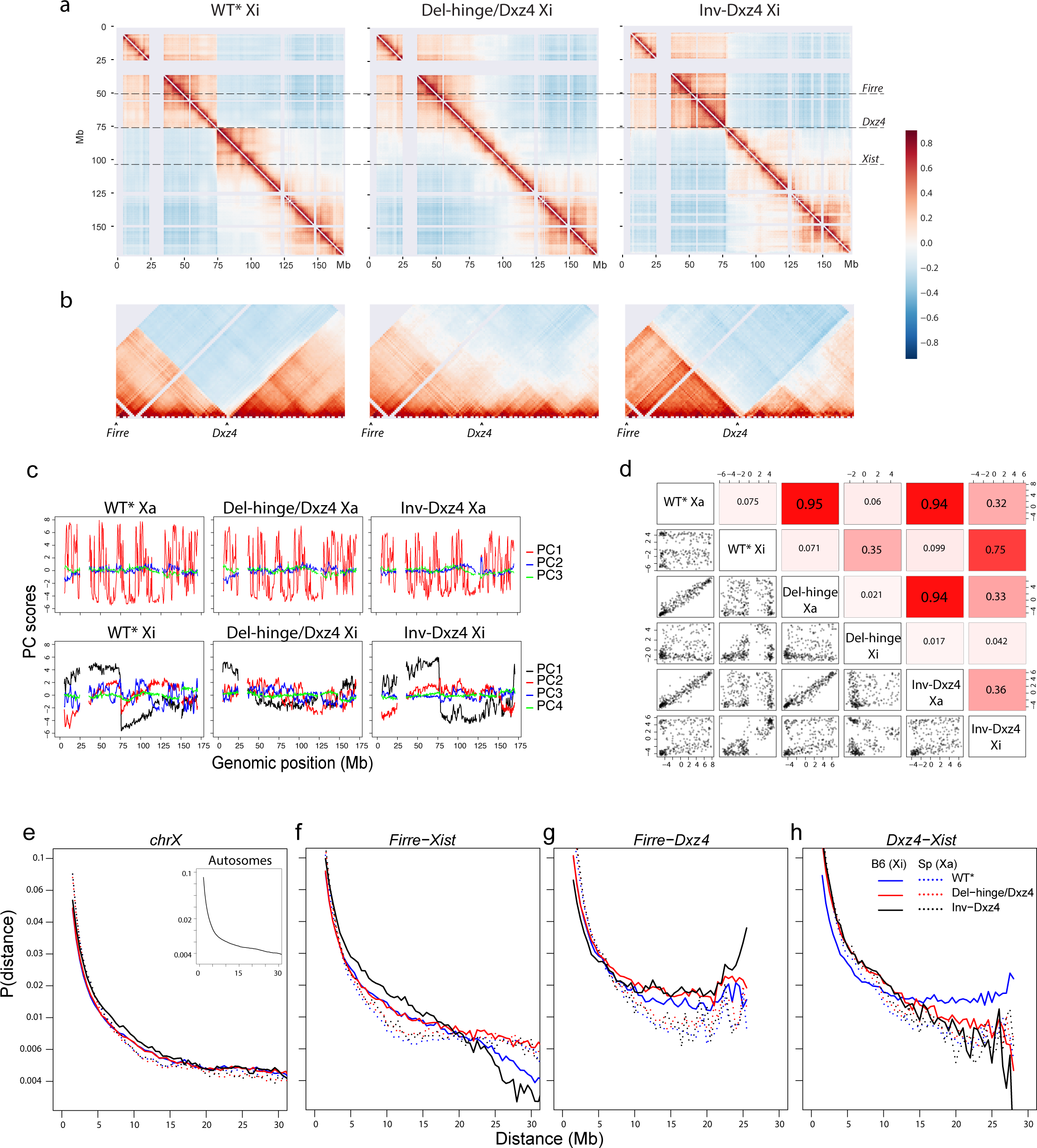
*Dxz4* deletion or inversion changes contact distribution on the Xi. **a**. Contact maps (500kb resolution) for the Xi in WT*, Del-hinge/Dxz4 and Inv-Dxz4 visualized using Pearson correlation to highlight contact probabilities. The color scale shows Pearson correlation values. **b**. Pearson correlation-transformed contact maps (500kb resolution) for 50Mb around the *Dxz4* locus to highlight the loss of superdomain structure in Del-hinge/Dxz4 and the shift in contacts in Inv-Dxz4, where a new contact domain forms between *Firre* and *Dxz4.* The color scale shows Pearson correlation values. **c**. Allelic principal component (PC) score profiles for the Xa and Xi chromosomes in WT*, Del-hinge/Dxz4, and Inv-Dxz4, based on distance-corrected, normalized contact maps with counts binned at 500kb resolution. For the Xa, the top three allelic PC scores are shown in red, blue and green, respectively. For the Xi, PC1 score profiles are plotted in black to emphasize that PC1 captures the bipartite structure rather than A/B compartments in WT*, and PC2-4 are shown in red, blue and green, respectively. See Supplementary Fig. S3a for analysis of the variance in the principal component score profiles of the X chromosomes and S3b-c for analyses of autosomes. **d**. Pairwise Spearman correlation values and associated scatterplots between allelic PC1 scores for the Xa and Xi in WT*, Del-hinge/Dxz4, and Inv-Dxz4. See Supplementary Fig. S3d for analysis of autosomes. **e-h**. Plots of the average Hi-C interaction frequencies (at 500kb resolution) as a function of genomic distance along the Xa (dotted line) and Xi (line) for WT* (blue), Del-hinge/Dxz4 (red) and Inv-Dxz4 (black) for the entire length of the X chromosomes (e) and for three regions along the X chromosome: *Firre-Xist* (~53Mb, f), *Firre-Dxz4* (~25Mb, g), *Dxz4-Xist* (~28Mb, h). Inset: plots of average contacts as a function of distance for autosomes.

To investigate whether disruptions of the Xi structure in Del-hinge/Dxz4 and Inv-Dxz4 result in changes to A/B compartment scores we decomposed WT*, Del-hinge/Dxz4, and Inv-Dxz4 allelic chromosomal contact maps into principal components (PCs). PC score profiles for autosomes and for the Xa were very similar between WT*, Del-hinge/Dxz4, and Inv-Dxz4, as expected, with PC1 capturing the A/B compartment structure (Fig. 2c; Supplementary Fig. S3a-c). In contrast, for the WT* Xi PC1, and to a lesser extent PC2 profiles capture the Xi bipartite structure rather than the underlying A/B compartment structure (Fig. 2c). Del-hinge/Dxz4 Xi PC1 score profile no longer shows the distinctive switch in sign at *Dxz4* seen in WT* reflecting the disruption to the bipartite structure (Fig. 2c). Indeed, WT* and Del-hinge/Dxz4 Xi PC1 scores show much lower correlation (Spearman *ρ* = 0.35) than seen for the Xa (Spearman *ρ* = 0.95) and autosomes (Spearman *ρ* > 0.9) (Fig. 2d; Supplementary Fig. S3d). Although the distinctive bipartite profile seen in the WT* Xi PC1 score is still evident for Inv-Dxz4 Xi PC1, their correlation is intermediate (Spearman *ρ* = 0.75), reflecting the changes to Inv-Dxz4 Xi long-range contacts (Fig. 2c, d).

To evaluate regional changes in contacts with respect to distance we generated contact decay curves representing the average contact count as a function of distance (see Methods). Plots of the contact decay curves for the whole X chromosome show largely similar trends to those for the autosomes, although the X chromosome plots also exhibit the expected differences between the active and inactive alleles (Fig. 2e). In contrast, strong differences between WT*, Del-hinge/Dxz4 and Inv-Dxz4 contact decay curves are observed at particular regions along the X chromosome. For instance, within the ~53Mb region from *Firre* to *Xist,* decay curves for Del-hinge/Dxz4 show a higher proportion of very long-range interactions (>20Mb) due to the loss of insulation between the two superdomains when *Dxz4* is absent. Indeed, the Del-hinge/Dxz4 profile is more similar to that of the Xa at these long distances (Fig. 2f). Conversely, within this same *Firre-Xist* region Inv-Dxz4 makes even fewer very long-range interactions (>20Mb) than WT*, implying that the inversion of *Dxz4* results in even stronger insulation between the two superdomains than that observed for the wild-type orientation (Fig. 2f). When only the region from *Firre* to *Dxz4* (~25Mb apart) is considered, both Del-hinge/Dxz4 and Inv-Dxz4 show a higher proportion of contacts than WT* within the 10–20 Mb range, with Inv-Dxz4 showing a higher proportion of very long-range contacts (>20Mb), reflecting the increase in contacts between the two loci (Fig. 2g). Contact decay curves for the region between *Dxz4* and *Xist* (~28Mb apart) highlight the dramatic loss in long-range contacts (>15Mb) between these two loci in the mutant lines compared to the wild-type (Fig. 2h). Note that a locus close to *Xist, X75* previously shown to interact with *Dxz4*^8^, may also contribute to these changes in contacts.

Differential contact maps confirm that the largest changes in contact frequency on the Xi in Del-hinge/Dxz4 or Inv-Dxz4 versus WT* are located in the vicinity of *Dxz4* and show unidirectionality (Fig 3a, b; Supplementary Fig. S3e-h). Contacts between the hinge region and loci telomeric to the locus (chrX:75-115Mb) decrease in both Del-hinge/Dxz4 and Inv-Dxz4, indicating that *Dxz4* normally contacts loci in the telomeric superdomain of the Xi. In Del-hinge/Dxz4 new long-range contacts appear between the two Xi superdomains, consistent with *Dxz4’s* role of insulating the two superdomains (Fig. 3a; Supplementary Fig. S3e, f). In Inv-Dxz4, new contacts appear between *Dxz4* and multiple loci in the centromeric superdomain (chrX:5-75Mb) (Fig. 3b; Supplementary Fig. S3g, h). Strikingly, a new contact domain appears between *Firre* and *Dxz4,* while the contact domain between *Dxz4* and *Xist* or *X75* present in WT* is lost (Fig. 3b; Supplementary Fig. S3g, h). These contact changes most probably result from the reversal of the orientation of CTCF motifs previously identified at *Dxz4,* which are mainly oriented toward the telomeric end of the mouse X in wild-type (Fig. 1b)^12^.

**Fig. 3:**
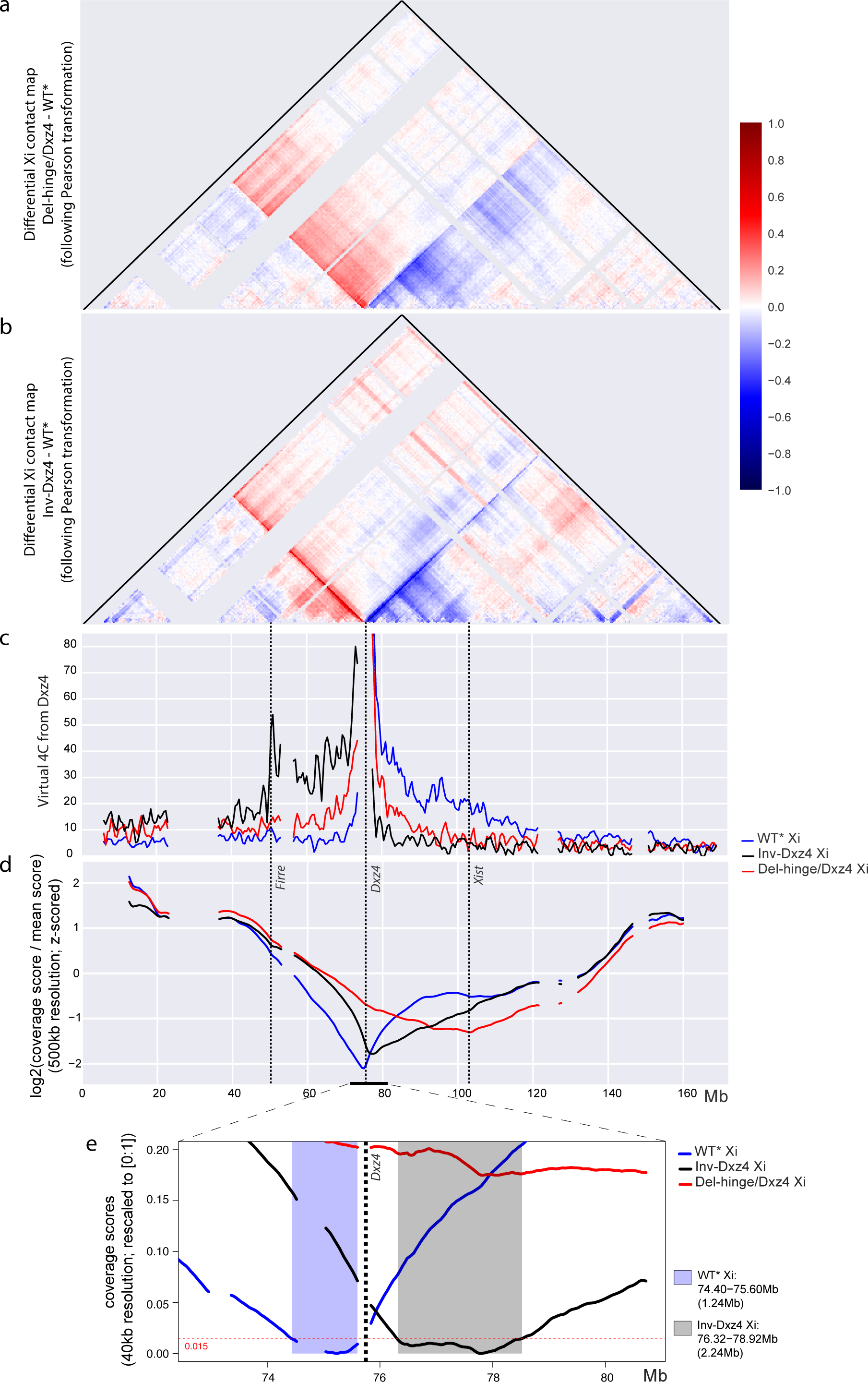
Unidirectional disruption of contacts on the Xi after *Dxz4* deletion or inversion. **a**. Differential contact map based on Pearson correlation transformed data at 500kb resolution to highlight differences between Del-hinge/Dxz4 Xi and WT* Xi (loss or gain of contacts in the Del-hinge/Dxz4 versus WT* appear blue or red, respectively). The color scale shows differential Pearson correlation values. See Supplementary Fig. S3e, f for comparison with differential contact maps based on untransformed count data. **b**. As in (a) to highlight differences between Inv-Dxz4 Xi and WT* Xi. See Supplementary Fig.S3g, h for comparison with differential contact maps based on untransformed count data. c. Virtual 4C plots derived from Hi-C data at 500kb resolution using a 500kb region around *Dxz4* as the viewpoint on the Xi in WT* (blue), Del-hinge/Dxz4 (red), and Inv-Dxz4 (black). Y-axis (contact counts) limited to 20% of maximum. The positions of *Firre, Dxz4* and *Xist* are indicated. See additional 4C analyses in Supplementary Fig. S3i. **d**. Standardized coverage score profiles at 500kb resolution for the Xi in WT* (blue), Del-hinge/Dxz4 (red), and Inv-Dxz4 (black). The positions of *Firre, Dxz4* and *Xist* are indicated. See additional analysis of coverage scores for Xi and Xa in all cell lines in Supplementary Fig. S4a-d. **e**. Coverage scores (rescaled to [0; 1]) at 40kb resolution within a 8Mb region around *Dxz4* for the Xi. The light blue background highlights the Xi boundary region of minimal interaction in WT* based on a threshold of 0.015 (horizontal red dashed line). The light grey background shows how this region is shifted to the right in Inv-Dxz4.

The unidirectional nature of interactions between *Dxz4* and other Xi loci is clearly evident in virtual 4C plots based on Hi-C data using *Dxz4* as a viewpoint (Fig. 3c). Additional 4C plots generated for additional viewpoints along the Xi show that interactions between *Dxz4* and regions located as far as 40Mb telomeric to the locus (chrX:75-115Mb) are strongly reduced in Del-hinge/Dxz4 or Inv-Dxz4 versus WT* (Supplementary Fig. S3i). Note that along this 40Mb region on the WT* Xi the number of contacts with *Dxz4* varies with evidence for contact hotspots, for example between *Dxz4* and either *Xist* or *X75* (Fig. 3c; Supplementary Fig. S3i). Contact hotspots were also seen in Inv-Dxz4, especially between *Dxz4* and *Firre* (Fig. 3c). To quantify changes in contact frequency extending across the hinge region at all scales, including very long-range contacts, we used a modified version of the coverage score measure^31^ (see Methods). The strong dip in coverage score due to lack of contacts between superdomains on the Xi in WT* was not seen in Del-hinge/Dxz4, but was mostly retained in Inv-Dxz4 (Fig. 3d). This analysis confirms extensive loss of long-range contacts between *Dxz4* and regions telomeric to the locus (chrX:75-115Mb) in Del-hinge/Dxz4, and gain of contacts between the two superdomains resulting in a higher coverage score at chrX:50-75Mb, while few changes occurred in distal regions (chrX:5-50Mb, 140-165Mb) (Fig. 3d). In the Inv-Dxz4 Xi, a dramatic decrease in the coverage score between chrX:75-110Mb and a corresponding increase between chrX:45-75Mb can be attributed to the reversal of *Dxz4* anchored contacts. Similar results were obtained when considering all cell lines: changes to the coverage score profile are only seen for the deletions that include *Dxz4* and the inversion of *Dxz4* (Supplementary Fig. S4a). Coverage scores for the Xa were similar between cell lines. Violin plots showed a smaller range in negative coverage scores for Del-hinge and Del-Dxz4, and to a lesser extent for Inv-Dxz4, than for WT, Del-Ds-TR, and Inv-5’ Ds-TR, confirming loss of very long-range contacts (Supplementary Fig. S4b). Hierarchical clustering using both Euclidean and Pearson correlation distance measures resulted in segregation of Xa’s from Xi’s, and within the Xi’s, segregation of those in Del-hinge and Del-Dxz4 from the others (Supplementary Fig. S4c, d).

Intriguingly, the differential contact map between Inv-Dxz4 and WT* shows that a region immediately telomeric to *Dxz4* makes many new contacts with centromeric loci in Inv-Dxz4 Xi. This phenomenon is visualized as a red band of differential contacts running from *Dxz4* to the centromere (Fig. 3b). Interactions with telomeric loci are simultaneously lost (blue band of differential contacts) from *Dxz4* to telomere. These bands represent the reversal of a line of long-range contacts, also called a “flame”^21,32^, emanating from *Dxz4* (Fig. 1a, 2b, 3b). This flame represents a shift in the transition zone between the two superdomains. Examining this region more closely using coverage scores based on higher resolution data (40kb), we found that the region of minimal interactions between the two superdomains shifts to the opposite side of *Dxz4* in Inv-Dxz4 Xi compared to WT*, while almost doubling in size (Fig. 3e).

Taken together, these results demonstrate that *Dxz4* specifically contacts a number of regions located telomeric to the locus in the wild-type Xi, and that deletion or inversion of *Dxz4* exerts their strongest effects in the vicinity of the locus. The unidirectional nature of contacts due to the orientation of CTCF binding sites is clearly demonstrated by contact changes observed after *Dxz4* inversion.

### 3. Xi TAD structure is largely unchanged following *Dxz4* deletion or inversion except around the *Dxz4* locus

We and others have reported that the mouse Xi does not display prominent TADs, in contrast to the Xa or the autosomes^7,9,10,19^. To quantify TAD number and distribution, we called TADs using the insulation score method^33^ at 500kb and 40kb resolution (Fig. 4a-c). Overall, a similar number of TADs were identified on the Xi and Xa in WT*; however, TAD insulation scores were attenuated on the Xi (Supplementary Table S3; Fig. S4e, f). Insulation scores for Del-hinge/Dxz4 and Inv-Dxz4 at 500kb resolution differed from those in WT*, especially around *Dxz4* (Fig. 4a; Supplementary Fig. S4e). This effect is more evident when considering the region between *Firre* and *Xist* using 40kb bins (Fig. 4b). The most distinctive change in this region is the loss of the deep trough at *Dxz4* when the locus is deleted, which results in the steep peak seen in the differential plot between Del-hinge/Dxz4 and WT* insulation scores (Fig. 4c). Xi insulation scores cluster away from those from the Xa, regardless of whether Euclidean or correlation (1 - *r*) distance metric is used (Fig. 4d, e;), and a similar allelic separation is seen when comparing the extent of TAD overlap (Fig. 4f). Additionally, boundary scores and insulation score amplitudes are much greater along the Xa than the Xi both chromosome-wide and within the region from *Firre* to *Xist* (Fig. 4g; Supplementary Fig. S4f). Clustering of the insulation scores obtained for each individual mutant did segregate scores for the Xi’s from Del-hinge, Del-Dxz4 and Inv-Dxz4 away from those of the samples with an intact bipartite structure (Del-Ds-TR, WT, Inv-5’Ds-TR), indicating that the mutants with a *Dxz4* deletion or inversion showed a similar subtle difference in their topological domain structures (Supplementary Fig. S4g).

**Fig. 4:**
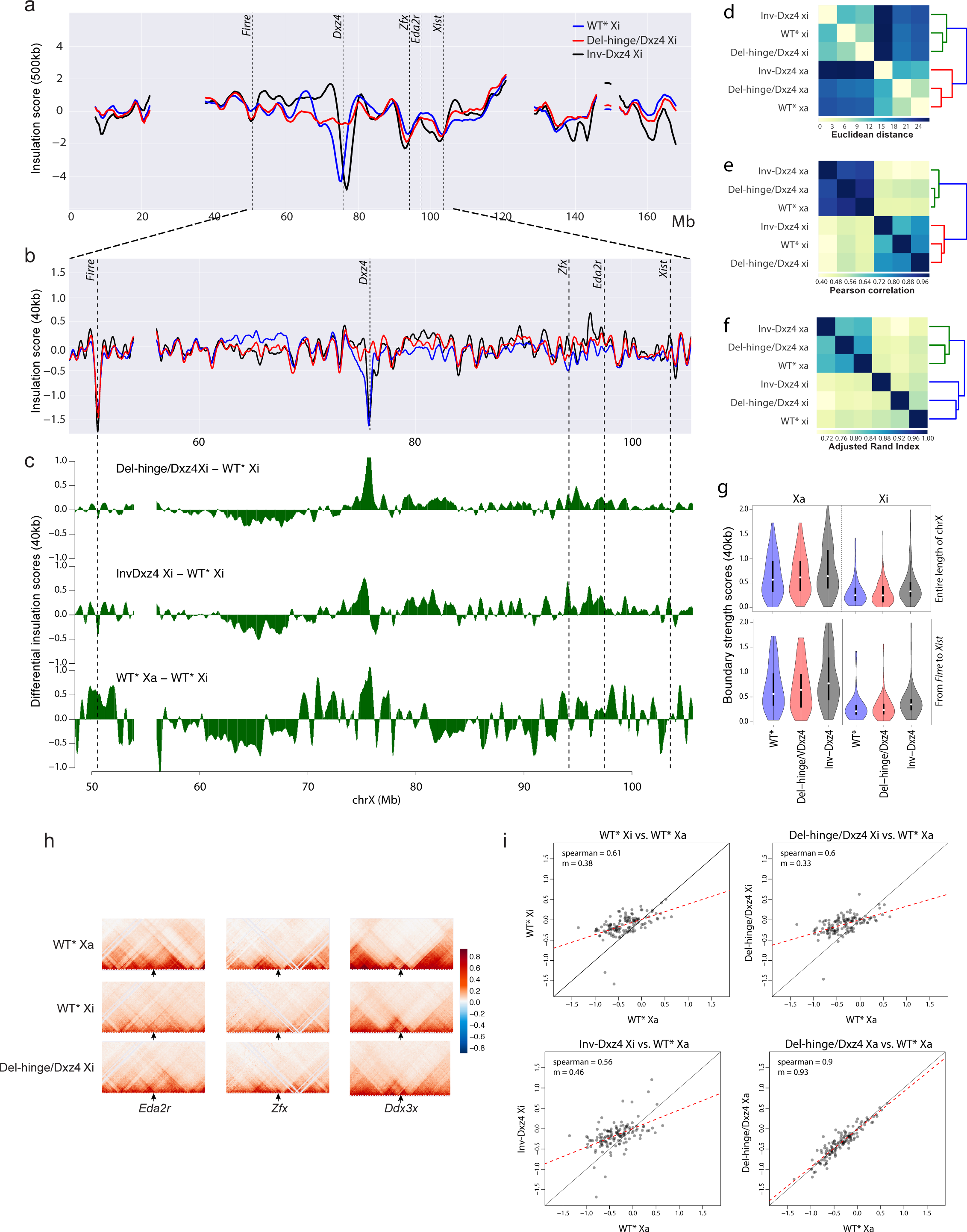
Changes in Xi TAD configuration after *Dxz4* deletion or inversion. **a**. Insulation score profiles at 500kb resolution for the whole Xi in WT* (blue), Del-hinge/Dxz4 (red), and Inv-Dxz4 (black). The positions of *Firre, Dxz4, Zfx, Eda2r* and *Xist* are indicated. See Supplementary Fig. S4e, f for analysis of insulation scores of Xi and Xa in all cell lines. **b**. As in (b) but based on 40kb resolution data for a region from *Firre* to *Xist* along the X chromosome. **c**. Differential insulation score profiles for Del-hinge/Dxz4 Xi (top), Inv-Dxz4 Xi (middle) and WT* Xa (bottom) relative to WT* Xi based on 40kb resolution data for a region from *Firre* to *Xist.* **d, e**. Hierarchical clustering based on the Euclidean distance (e) and Pearson correlation (using 1 - *r* as the distance measure; f) between standardized insulation scores at 40kb resolution considering the Xi and Xa in WT*, Del-hinge/Dxz4 and Inv-Dxz4. See Supplementary Fig. S4g for analysis of all cell lines. **f**. Hierarchical clustering based on the adjusted Rand index to quantify the correspondence between TADs called using insulation scores at 40kb resolution along the Xi and Xa in WT*, Del-hinge/Dxz4 and Inv-Dxz4. **g.** TAD boundary strength scores (see methods) for the Xa (left) and Xi (right) in WT*, Del-hinge/Dxz4 and Inv-Dxz4 for the entire X chromosome (top) and for the region from *Firre* to *Xist* (bottom). **h**. Comparisons of contacts on the Xa and Xi in WT* and Del-hinge/Dxz4 at individual loci. Changes in TAD configuration at 40kb resolution are shown within 4Mb regions, each centered at a specific gene, including two genes normally subject to XCI, *Edar2* and *Zfx,* and a gene that escapes XCI, *Ddx3x.* **i**. Scatter plots of WT* Xi, Del-hinge/Dxz4 Xi, Inv-Dxz4 Xi and Del-hinge/Dxz4 Xa insulation scores at the TAD boundaries identified on WT* Xa.

Inversion of *Dxz4* also results in a dramatic change in insulation score immediately around the locus, with a rapid switch in the differential plot corresponding with a shift of the *Dxz4* trough towards the telomeric end of the Xi (Fig. 4a-c; Supplementary Fig. S4e, f). Both Del-hinge/Dxz4 and Inv-Dxz4 show an increase in short to medium range contacts in a region 5-10Mb telomeric to *Dxz4,* as well as around *Zfx* and *Eda2r,* consistent with restoration of TADs (Fig. 4b, c; Supplementary Fig. S4f). Zooming in to assess potential changes in short-range contacts around loci normally subject to XCI (e.g. *Eda2r, Zfx),* shows that TADs, normally attenuated on the Xi, are partially restored on the deleted Xi in Del-hinge/Dxz4 and assumes a pattern more similar to that of the Xa (Fig. 4h). In contrast, loci that escape XCI, such as *Ddx3x,* show little change. A particularly large decrease in Del-hinge/Dxz4 and Inv-Dxz4 Xi insulation scores relative to WT* Xi occurs in a region between 60 and 68Mb along the Xi, a region where WT* Xa insulation is also lower than WT* Xi (Fig. 4b, c; Supplementary Fig. S4f). Scatter plots of Xi insulation scores at WT* Xa boundary regions (using 40kb bins) show that the Del-hinge/Dxz4 and Inv-Dxz4 Xi insulation scores at these positions display a similar trend to that observed for WT* Xi, whereas similar plots for the Xa in Del-hinge/Dxz4 and Inv-Dxz4 show a much closer correlation with insulation scores at WT* Xa boundaries (Fig. 4i and data not shown). We conclude that the Xi TAD structure remains largely unchanged along the Xi, except at select loci and around the *Dxz4* locus following its ablation or inversion.

### 4. CTCF binding to the Xi increases after *Dxz4* deletion or inversion

CTCF plays an important role in defining boundaries between contact domains, such as TADs^18-20^. CTCF-ChIP analyses confirmed loss of CTCF binding at the deleted *Dxz4* locus in Del-hinge cells,while CTCF binding at the imprinting control region near *H19* was maintained (Supplementary Fig. S5a, b). We derived CTCF binding profiles by ChIP-seq for WT, Del-hinge and Inv-Dxz4 Patski cells calling CTCF peaks jointly across both alleles using unsegregated reads (Supplementary Table S4). The majority of CTCF peaks contained SNPs and were covered by at least five SNP reads (Supplementary Fig. S5c-e). For each CTCF peak with sufficient SNP coverage (5x), an allelic proportion of SNP read coverage (*spretus*/(*spretus*+BL6)) was calculated. Average allelic proportions (Xa/(Xa+Xi)) of X-linked CTCF peaks are markedly different between both Del-hinge (~60%) and Inv-Dxz4 (~63%) compared to WT (~71%) while average allelic proportions for autosomes are similarly close to 50% in all samples as expected (Supplementary Fig. S5f). This difference is also reflected in the distributions of allelic proportion for CTCF peaks along the X chromosome, which show a pronounced shift toward lower values in Del-hinge and Inv-Dxz4 compared to WT (with modes of 0.57, 0.6, and 0.84, respectively) (Fig. 5a; Supplementary Fig. S5g). In addition, analysis of Patski2-4, a wild-type subclone used to derive the inversion line showed similar results to the original WT cell population with CTCF peaks along the X chromosome showing an average allelic proportion of 67% (data not shown). Plots of CTCF peak d-scores (*spretus*/(*spretus*+BL6) - 0.5) confirm the increase in CTCF binding along the entire Xi in Del-hinge and to a lesser extent in Inv-Dxz4 whereas d-scores fall mainly around 0.5 along autosomes (chromosome 2 for example) (Supplementary Fig. S6a). Imprinting control regions near *H19* (maternally expressed gene) and within *Peg3* (paternally expressed gene) on chromosome 7 showed CTCF peaks only on the maternal and paternal allele, respectively (Supplementary Fig. S6a).

**Fig. 5.**
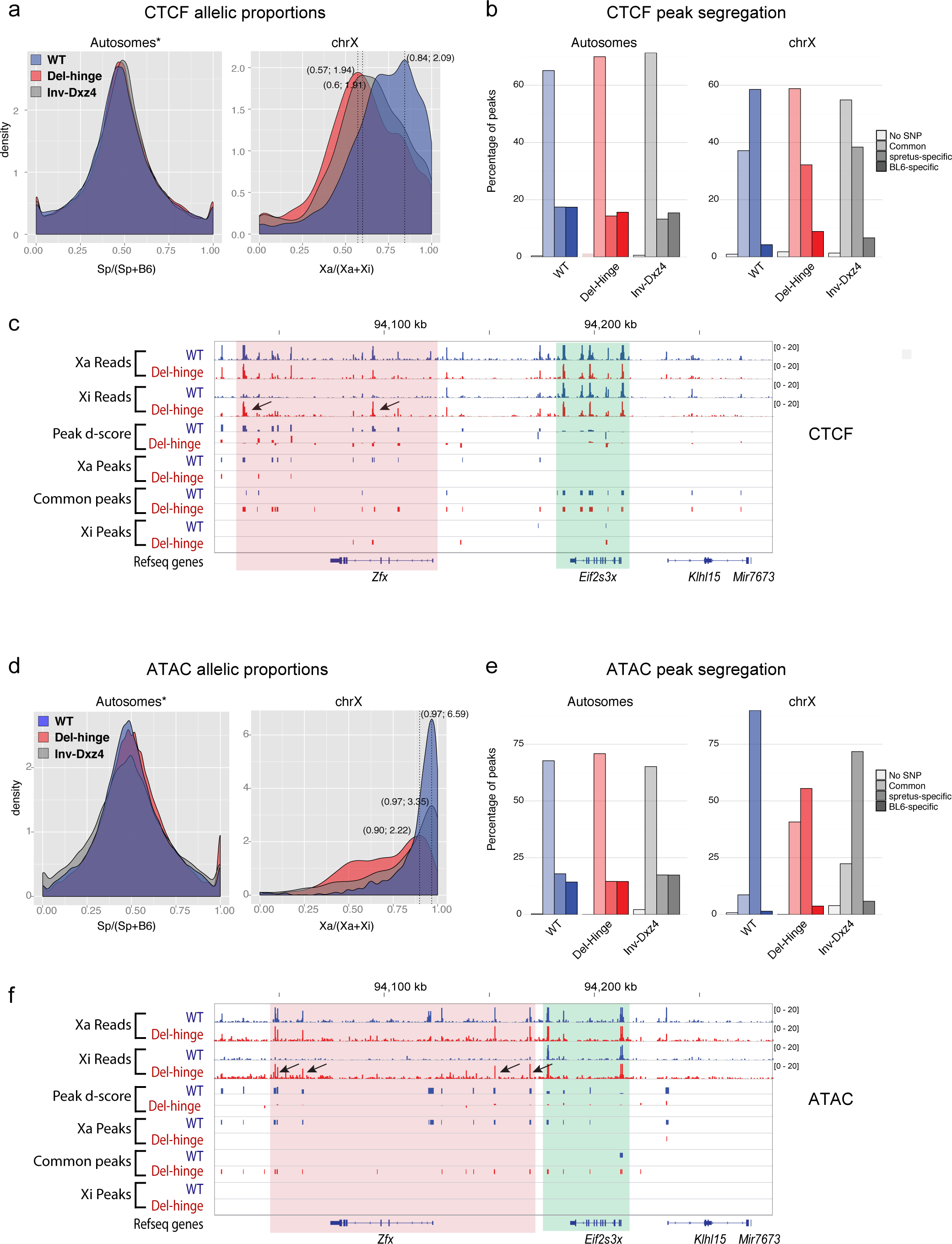
Xi CTCF and ATAC peak distribution changes in Del-hinge and Inv-Dxz4 cells. **a**. Density histograms of the distribution of allelic proportions of CTCF peaks *(spretus/(spretus* +BL6) along autosomes and the X-chromosomes for WT (blue), Del-hinge (red) and Inv-Dxz4 (grey). The modes of the X-chromosome distributions are given in parentheses. *Chromosomes 3 and 4 were removed from the autosomes because they show aneuploidy. See also Supplementary Fig. S5g. **b**. Percentages of CTCF peaks in WT (blue) Del-hinge (red) and Inv-Dxz4 along the autosomes and the X-chromosomes classified as spretus-specific, BL6-specific, or common peaks. **c.** Genome browser tracks of CTCF peaks on the Xa and Xi, of CTCF peak d-scores ((Xa/(Xa+Xi) - 0.5), and of CTCF peaks assigned as Xa-specific, common, or Xi-specific for WT (blue), Del-hinge (red) and Inv-Dxz4 along a region of the X-chromosome that includes *Zfx* (a gene subject to XCI in WT, which reactivated in Del-hinge) and *Eif2s3x* (a gene that escapes XCI). CTCF peaks that appear on the Xi in a region around *Zfx* in Del-hinge are indicated with arrows. **d-f**. As in a-c but for ATAC peaks. See additional analyses in Supplementary Fig. S5-S7.

Diploid CTCF peaks with sufficient SNP coverage (5x) were designated as either *spretus*-specific or BL6-specific based on an allelic proportion (spretus/(spretus+BL6)) greater than 70% or lower than 30%, respectively, with remaining peaks classified as common (Fig. 5b). The majority of X-linked CTCF peaks in Del-hinge and Inv-Dxz4 are common (~59% and ~55%, respectively), whereas in WT there is a comparatively lower proportion of common peaks (37%) with the majority being Xa-specific (>58%), reflecting an increase in CTCF binding along the Xi after deletion of the hinge or inversion of *Dxz4* (Fig. 5b). By comparison, the majority of autosomal peaks are common in all three cell lines (65-71%). The additional common CTCF peaks seen for Del-hinge and Inv-Dxz4 appear to be found within similar regions all along the X chromosome to where WT common CTCF peaks bind such that the binned counts correlate very strongly, with these regions corresponding to regions where Xa-specific peaks also tend to be found (Supplementary Fig. S6c). Although far fewer in number, the same appears to be true for Xi-specific peaks. We conclude that deletion of the hinge and to a lesser extent inversion of *Dxz4* result in a general increase in CTCF binding along the Xi. As described below, neither *Xist* expression nor *Xist* RNA coating of the Xi were measurably altered in the mutant cell lines, suggesting that the changes in CTCF binding observed are mostly independent of *Xist.*

Interestingly, some of the CTCF peaks that appear on the Xi in Del-hinge cells occur near genes whose TAD structure and expression change. For example, at *Zfx,* a gene normally subject to XCI, new CTCF binding peaks (marked by arrows) located both within the gene and downstream of the gene appear on the Del-hinge Xi, consistent with more prominent TADs as well as reactivation of the gene (see below) (Fig. 4h; 5c). In contrast, genes that escape X inactivation both in WT and Del-hinge cells, for example *Eif2s3x,* show no change in CTCF peaks (Fig. 5c).

### 5. Chromatin accessibility increases on the Xi after deletion or inversion of the hinge

Chromatin accessibility was measured using ATAC-seq. As was done for CTCF, for both WT, Del-hinge and Inv-Dxz4 ATAC peaks were called jointly across both alleles using unsegregated reads (Supplementary Table S5). The number of SNP-containing peaks was similar between the three cell lines, and the vast majority of SNP-containing ATAC peaks were covered by at least 5 reads (Supplementary Fig. S7a-c). Although the overall autosomal allelic proportions (*spretus*/(*spretus*+BL6)) were as expected (~50%), the X-chromosome allelic proportion for Del-hinge, and to a lesser extent for Inv-Dxz4, were much lower compared to WT (~70% and 78% compared to ~88%), indicative of an increase in ATAC peaks and chromatin accessibility in the mutant cell lines (Supplementary Fig. S7d). Indeed, the WT, Del-hinge and Inv-Dxz4 distributions of allelic proportion for ATAC peaks are shifted for the X-chromosome, while for autosomes the distributions overlap almost perfectly (Fig. 5d; Supplementary Fig. S7e). Plots of ATAC peak d-scores (*spretus*/(*spretus*+BL6) - 0.5) show that the increase in chromatin accessibility occurs across the entire Xi in Del-hinge and Inv-Dxz4, rather than being restricted to regions around *Dxz4* (Supplementary Fig. S6b). Autosomes, on the other hand, show little to no difference in d-score between WT and mutant cell lines.

Diploid ATAC peaks with sufficient SNP coverage (5x) were designated as *spretus*-specific or BL6-specific if they showed an allelic proportion (*spretus*/(*spretus*+BL6)) greater than 70% or lower than 30%, respectively, and common otherwise (Fig. 5e; Supplemental Fig. S7e). A far smaller proportion of ATAC peaks are Xa-specific in Del-hinge (~55%) compared to WT (~90%), with a far higher proportion of common peaks in Del-hinge (40%) compared to WT (~9%) (Fig. 5e). In Inv-Dxz4 these proportions are intermediate between Del-hinge and WT with ~72% peaks being Xa-specific and 23% common to both alleles. In addition, analysis of Patski2-4, a wild-type subclone used to derive the inversion line, showed similar results to the original WT with ~90% peaks being Xa-specific and only ~7% common (data not shown). In contrast, autosomal ATAC peaks show comparable proportions of common and allele-specific peaks, implying similar accessibility in all three cell lines (Fig. 5e). Genome browser tracks clearly show a strong increase in ATAC peaks along the Xi (BL6 X-chromosome) in Del-hinge and Inv-Dxz4 cells, which is something not seen along BL6 autosomes (Supplementary Fig. S6b). Zooming in on individual genes, the appearance of ATAC peaks around X-linked genes that are normally inactivated and become expressed from the Xi in Del-hinge cells, for example *Zfx,* is evident, while ATAC peaks are not altered around genes that escape XCI such as *Eif2s3x* (Fig. 5f).

As seen for CTCF peaks, the additional Del-hinge and Inv-Dxz4 common ATAC peaks are found within similar regions along the X chromosome to WT common and Xa-specific ATAC peaks (Supplementary Fig. S6d). The same appears to be true for Xi-specific peaks, although the correlation is lower, presumably due their relative sparsity. Furthermore, binned CTCF and ATAC common peaks correlate both in Del-hinge (Spearman *ρ* = 0.84) and Inv-Dxz4 (Spearman *ρ* = 0.6), indicating the correlation of these two features. Indeed, unbinned common CTCF peaks and ATAC peaks showed spatial colocalization by the relative distance metric^34^ along both the Xa and Xi in WT, Del-hinge and Inv-Dxz4, which is also the case for Xa- and Xi-specific peaks (data not shown). Taken together, results of the ATAC-seq analysis show a strong increase in Xi chromatin accessibility after deletion of the hinge and a lesser increase after inversion of *Dxz4,* consistent with a role for *Dxz4* in maintaining condensation of the Xi.

### 6. Xi volume, *Xist* RNA coating, H3K27me3 accumulation and Xi positioning after *Dxz4* deletion or inversion

To test for changes in the 3D volume of the Xi in cell lines with deletion or inversion of *Dxz4* we examined interphase nuclei by fluorescence microscopy after DNA-FISH using a whole X chromosome paint together with H3K27me3 immunostaining to locate the Xi (Fig. 6a). The fluorescence intensity of signals for the X paint, H3K27me3, and Hoechst staining were comparable between wild-type Patski2-4, Del-hinge and Inv-Dxz4 on both the Xi and Xa (Fig. 6b-d). In particular, H3K27me3 immunostaining resulted in a strong Xi-staining cluster in ~90% of the nuclei from WT, Del-hinge and Inv-Dxz4 cells (Fig. 6a, d), suggesting that *Dxz4* deletion or inversion did not cause major changes in enrichment of the repressive histone mark on the Xi. In contrast, image analyses showed a small increase in the volume of the Xi normalized to that of the Xa in Del-hinge (7%, *p*-value = 0.01 using Wilcoxon sum rank test) and Inv-Dxz4 (6%, *p*-value = 0.03) nuclei compared to wild-type Patski2-4 nuclei (Fig. 6e). RNA-FISH for *Xist* showed no difference in the percentage of nuclei with an *Xist* RNA cluster seen in WT (97%), Patski2-4 (91%), Del-hinge (95%) and Inv-Dxz4 (94%) cells, indicating that deletion or inversion of *Dxz4* did not disrupt *Xist* RNA coating on the Xi (Fig. 6f). Consistently, *Xist* expression levels were similar in WT, Patski2-4, Del-hinge and Inv-Dxz4 cells as shown by RNA-seq and quantitative RT-PCR (Supplementary Fig. S8a, b). Taken together, these results show that the changes in the configuration of the Xi we detect by Hi-C in Del-hinge/Dxz4 and in Inv-Dxz4 are associated with a small increase in the apparent size of the Xi, but not with any apparent changes in H3K27me3 accumulation or *Xist* RNA coating on the Xi. We cannot exclude changes that may be visible by super-resolution microscopy.

**Fig. 6.**
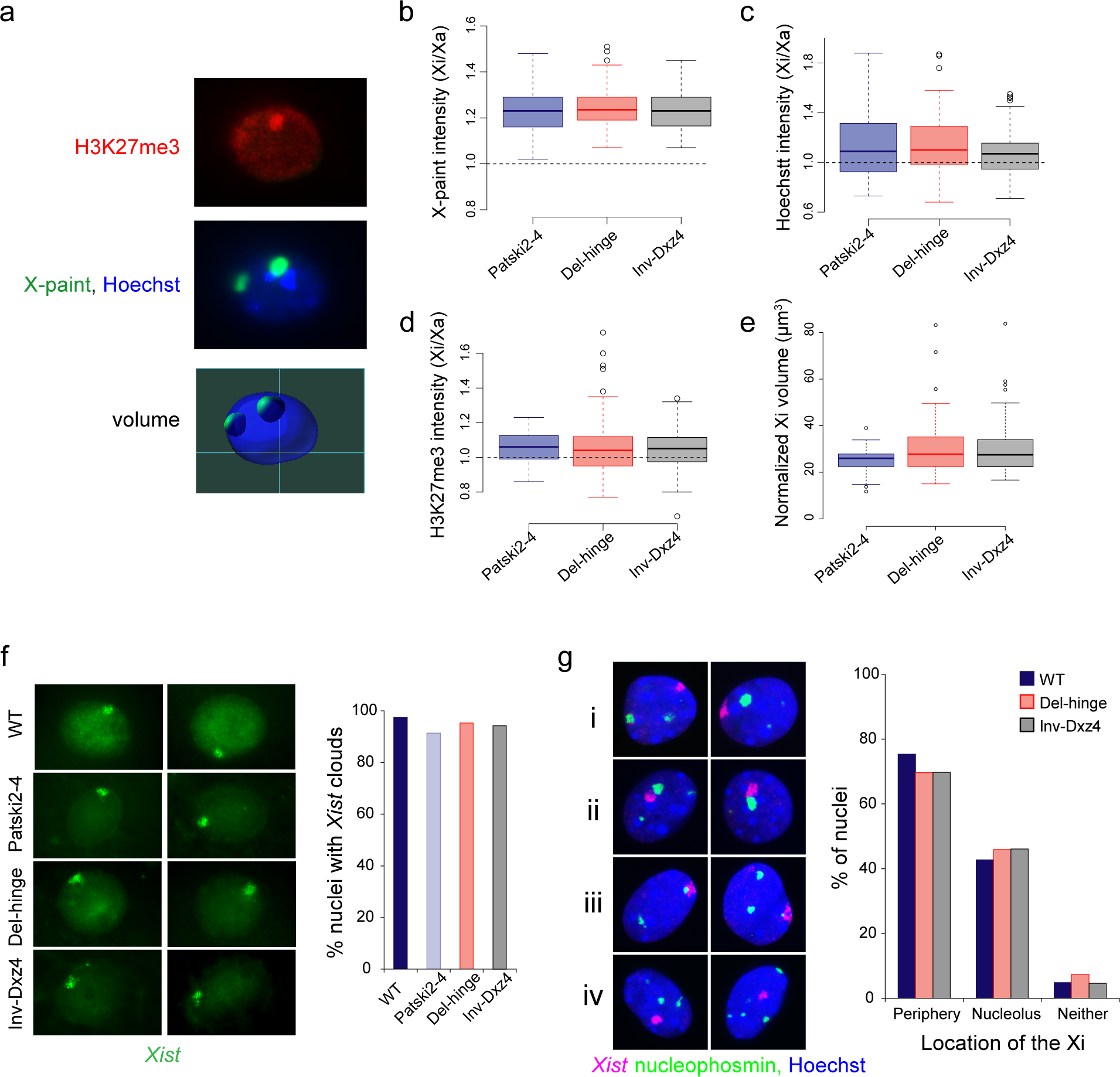
3D Xi volume, *Xist* RNA coating, H3K27me3 accumulation and Xi location after *Dxz4* deletion or inversion. **a**. Example of a z-stack image of Del-hinge nuclei after DNA FISH with H3K27me3 immunostaining (red), an X-paint probe (green), and DNA staining with Hoechst (blue). The volume of each X chromosome was called based on X-paint probe (the lines show the grid used in the microscope). The Xi was identified as that enriched in H3K27me. **b-d**. Box plots for X-paint signal intensity, Hoechst staining intensity and H3K27me3 immunostaining intensity between wild-type Patski2-4, Del-hinge and Inv-Dxz4 cells. No difference was detected between these lines by Wilcoxon rank sum test. Note that the Xi shows higher intensity of staining compared to the Xa. **e**. Box plots of the Xi volume normalized to the Xa volume show a slight but significant by Wilcoxon rank sum test (see text) increase of the median value in the Xi volume in Del-hinge (7%) and in Inv-Dxz4 (6%). **f**. Examples of nuclei from WT, Patski2-4, Del-hinge and Inv-Dxz4 cells after RNA-FISH for *Xist* (green) alongside bar plots of the percentage of nuclei with *Xist* RNA clouds. No significant difference was detected between cell lines by Fisher’s exact test. See Supplementary Fig. S8a, b for *Xist* expression analyses. **g**. Left, examples of nuclei stained with Hoechst (blue) and immunostained using an H3K27me3 antibody to locate the Xi (red), and an antibody to nucleophosmin to locate the nucleolus (green). The Xi was either located at the periphery (i), near the nucleolus (ii), or sandwiched between the periphery and the nucleolus (iii), or close neither to the periphery nor to the nucleolus (iv). Right, percentage of nuclei with the Xi near the periphery, the nucleolus, and neither of these locations in WT, Del-hinge and Inv-Dxz4. For each category no significant difference was detected between these cell lines by Fisher’s exact test.

We previously reported that the *Dxz4* and *Firre* loci on the Xi, but not on the Xa, are often preferentially located near the edge of the nucleolus, and that knockdown of *Firre* lncRNA caused a significant reduction in both the frequency of this preferred location and in the level of H3K27me3 on the Xi^15^. To determine whether *Dxz4* deletion or inversion affects Xi positioning within the nuclei of Patski cells, we performed dual immunostaining for H3K27me3, to mark the Xi, and for nucleophosmin, to label the nucleolus (Fig. 6g). No significant changes in the frequency of Xi positioning either near the nuclear periphery or near the nucleolus were found (Fig. 6g). Thus, neither *Dxz4* deletion, nor its inversion, affect the position of the Xi in the nucleus of Patski cells.

### 7. Gene expression after deletion or inversion of *Dxz4*

RNA-seq was performed in wild-type cells (WT and Patski2-4), in Del-hinge cell clones a and b, in Inv-Dxz4 clones a and b, and in single clones for Del-Dxz4 and Del-Ds-TR. An allele-specific data analysis pipeline (see Methods) identified a similar set of genes that escape XCI in WT cells as in our previous study (29 escape genes) (Supplementary Table S7)^35^. We initially focused our analyses on Del-hinge clone a, which is derived from WT cells. There was some evidence of reactivation of a few genes on the Xi, with 16 genes showing a significant increase in expression relative to WT, and no gene showing significant downregulation (log2 fold change > 0.5 and adjusted *p*-value < 0.05) (Fig. 7a; Supplementary Fig. S8c, Table S8). This unidirectional trend was not observed for the Xa, for which 55 genes showed a significant change in expression between Del-hinge and WT but with the change occurring in both directions (Supplementary Fig. S8c, Table S8). A total of 2689 autosomal genes showed a significant increase or decrease in gene expression (Supplementary Table S9). When X-linked genes were grouped in terms of their silencing/escape status in WT cells, we found that deletion of the hinge affected Xi-expression of a greater number (but lower proportion) of genes subject to XCI (12/331 or 4% of genes) than escape genes (4/29 or 14% of genes). Interestingly, reactivated genes are distributed all along the Xi in regions of high gene density, with no correlation between the position of *Dxz4* and the location of Xi-alleles with expression changes (Fig. 7b). Intriguingly, both the genes whose surrounding TAD structure becomes more similar to that of the Xa in Del-hinge (*Zfx, Eda2r*) also show significant Xi-specific upregulation (Fig. 4h, Table S8). Furthermore, in the case of *Eda2r* there appeared to be an associated increase in accessibility and CTCF binding in the surrounding region along the Xi in Del-hinge (Fig. 5 c, f)

**Fig. 7.**
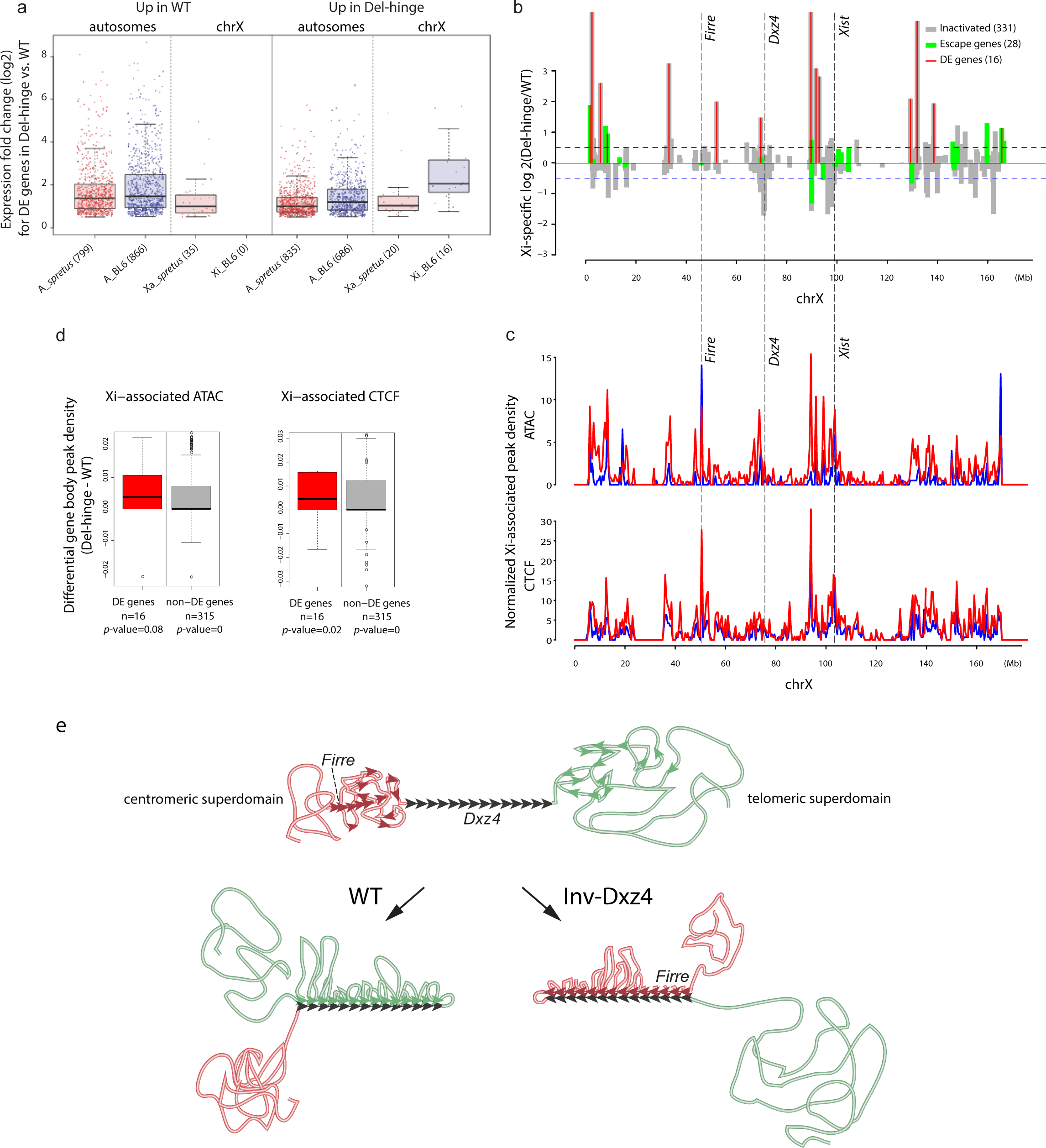
Gene expression changes after deletion of the hinge and model for the role of *Dxz4* in shaping the Xi. **a**. Box plots showing the distribution of fold changes in allelic gene expression in Del-hinge versus WT for autosomal (A_spretus or A_BL6) and X-linked (Xa or Xi) genes that have increased expression in WT or in Del-hinge. Red, *spretus* alleles; blue, BL6 alleles. **b**. Distribution of genes with expression fold changes between Del-hinge versus WT along the length of the Xi, considering 331 genes normally subject to XCI (grey) and 28 genes that escape XCI (green) (Note only 28/29 escape genes could be tested for differential expression because *Slc16a2* lacked the allelic coverage to do so). The 16 genes that show significant differential expression are demarcated with red vertical lines. The positions of *Firre, Dxz4, and Xist* are indicated. **c**. Plots of Xi-associated (common + Xi-specific) ATAC and CTCF peak density (counts binned within 500kb windows) along the X chromosome for WT (blue) and Del-hinge (red) aligned with expression fold changes in (a). To account for differences in the number peaks obtained between samples, the binned counts are scaled by a factor obtained from the between-sample ratios of total diploid autosomal peaks. For the ATAC peaks, the scaling factors are 0.62 and 0.63, while for CTCF the scaling factors are 0.7 and 0.78 for WT and Del-hinge, respectively. The positions of *Firre, Dxz4,* and *Xist* are indicated. See also Supplementary Fig. S6c, d and S8c. **d**. Boxplots of differential Xi-associated ATAC and CTCF peak densities between Del-hinge and WT. Peak densities within X-linked gene promoter regions and gene bodies (TSS - 10kb to TTS) are normalized for gene length and scaled by total diploid autosomal peak ratios. Genes are partitioned into those that show significant differential expression between WT and Del-hinge (DE; red) and those that did not (non-DE; grey). The number of genes in each group are shown. The *p*-values from a one-sided (>) paired Wilcoxon tests are also given. **e**. Models for the role of *Dxz4* in establishing long-range contacts with other loci. Diagram of the *Dxz4* locus (black) with adjacent centromeric (red) and telomeric (green) superdomains of the Xi. The orientation of CTCF binding motifs is shown with black arrows, with 14 motifs being represented. Potential CTCF sites located in the telomeric superdomain are shown as green arrows, and on the centromeric domain as red arrows. In WT cells contacts between *Dxz4* and loci telomeric to the locus would result in the formation of loops anchored at *Dxz4* by the correct alignment of CTCF motifs, which would stall cohesin rings that continuously extrude loops (not depicted). *Dxz4* would be pulled to the telomeric end of the hinge. After *Dxz4* inversion contacts would shift to the centromeric superdomain, and be especially enhanced between *Dxz4* and *Firre.* A new de-condensed hinge would form, with *Dxz4* located at its centromeric end. Note that in WT and Inv-Dxz4, loops are depicted as anchored at each CTCF binding site on *Dxz4.* However, the process of loop formation is not static, but rather highly dynamic; thus at a given time some of the loops would not be anchored and larger or smaller loops may form. In addition, one CTCF molecule at *Dxz4* may be sufficient to stall cohesin, instead of two as depicted here.

To determine whether there was a correlation between increased expression and increased accessibility and CTCF binding in general, we compared the location of genes that saw significant expression changes in Del-hinge clone a to the profiles of ATAC and CTCF peak density (Xi-specific and common peaks binned within 500kb) obtained in the same clone (Fig. 7b, c). The location of reactivated genes (i.e. those that significantly increased in Del-hinge versus WT) on the Xi fall within regions with a higher density of Xi-associated ATAC and CTCF peaks in Del-hinge, suggesting that decondensation of the Xi facilitates reactivation (Fig. 7b, c; Supplementary Fig. S6c, d). To determine whether genes upregulated along the Del-hinge Xi are particularly enriched for Xi-associated increases in accessibility and CTCF-binding, we calculated the ATAC and CTCF peak densities within all X-linked gene bodies and their upstream promoter regions (TSS - 10kb to TSS), with the genes grouped into those that showed significant differential expression between WT and Del-hinge (DE) and those that did not (non-DE) (Fig. 7d). Both ATAC and CTCF peak density were greater in Del-hinge for the 16 DE-genes, though this trend was only significant for CTCF binding (*p*-value = 0.02) and not for accessibility (p-value = 0.08). Non-DE genes also saw significant increases in accessibility (*p*-value = 0.02) and CTCF binding (p-value < 2e-16) in Del-hinge (Fig. 7d), indicating that although ATAC and CTCF densities along the Xi in Del-hinge are strongly correlated (Spearman *ρ* = 0.71), they do not correlate with the upregulation of any X-linked gene in particular but rather with gene dense regions (Spearman *ρ* = 0.41 and 0.57 for ATAC and CTCF, respectively). We conclude that the overall de-condensation and increased accessibility of the Del-hinge Xi does not result in the specific upregulation of Xi-linked genes, but the changes to the chromosome structure may set up an environment that allows for limited and stochastic reactivation/upregulation of genes.

Next, we compared Xi- and Xa-specific expression levels in additional cell lines including Del-hinge clone b, Del-Dxz4, and Del-Ds-TR to levels in WT (Supplementary Fig. S9a-d). As many as 77-85% of genes showed no or very low Xi-specific expression (Xi-TPM <0.2), indicating that silencing of most X-linked genes is largely maintained in these cell lines. There was a minor reactivation of the Xi in Del-hinge clone b and in Del-Dxz4, with the upregulated genes partially overlapping those observed in Del-hinge clone a, indicating some similarity but also variability between cell clones (Supplementary Fig. S9a-c, Table S8). This again suggests stochastic reactivation of genes in different cell clones. RNA-seq performed in two Inv-Dxz4 clones (a and b) and the corresponding subclones of wild-type cells used to derive the inversion lines (Patski2-4 a and b, respectively) showed no evidence of reactivation of genes on the Xi (log2 fold change > 0.5 and adjusted *p*-value < 0.05) (Supplementary Fig. S8d, S9e, Table S10). Only two genes (*Msn* and *Clcn5*) showed a significant increase in Xi-specific expression relative to wild-type, with another two genes (*Shroom2* and *Hnrnph2*) showing a significant decrease in Xi expression. By RT-PCR, *Dxz4* expression was similar in Del-hinge, Inv-Dxz4 and WT (data not shown), but we could not separate expression from the Xa or Xi, due to lack of informative SNPs.

XCI is associated with multiple layers of epigenetic modifications, including DNA methylation at CpG islands of silenced genes^1,2^. To test for potential synergistic effects between condensation of the Xi and DNA methylation, we examined the effects of global DNA demethylation on X-linked gene expression in Del-hinge clone a versus WT cells. Consistent with the inhibitory role of DNA methylation on gene expression, RNA-seq analyses showed global upregulation of both autosomal and X-linked genes in WT and Del-hinge cells after 5-aza-2dC treatment (Supplementary Fig. S10a-c). In WT cells, 2328 autosomal and 129 X-linked genes were significantly upregulated, with only 640 autosomal and 27 X-linked genes being downregulated (log2 fold change >0.5, adjusted *p*-value <0.05) (Supplementary Fig. S10a-c). Similarly, in Del-hinge cells, 1318 autosomal and 75 X-linked genes were significantly upregulated and only 133 autosomal genes and no X-linked gene, downregulated. Demethylation caused an increase in the number of reactivated X-linked genes, which was similar in Del-hinge versus WT (Supplementary Fig. S10a-c, Table S11, S12). Thus, deletion of *Dxz4* or of the hinge with or without DNA demethylation and inversion of *Dxz4* are insufficient to cause massive reactivation of genes on the Xi, despite the observed de-condensation and increase in chromatin accessibility in the mutant cell lines.

## Discussion

We determined that *Dxz4* is the sole element necessary for maintenance of the condensed 3D bipartite configuration of the inactive mouse X chromosome. Two previous studies based on large deletions (200-300kb) obtained in mouse ES cells or in human cells also reported disruption of the Xi bipartite structure^8,9^. However, we find that a much smaller deletion (44kb) of *Dxz4* alone is sufficient to cause de-condensation of the Xi. We found that both deletion of the hinge or of *Dxz4* cause few changes in the Xi TAD structure, except in the region around *Dxz4* and at individual loci. Remarkably, the Xi bipartite structure persists after inversion of *Dxz4,* indicating that the locus insulates the Xi superdomains in either orientation. However, we observed a massive redistribution of long-range contacts from the telomeric to centromeric superdomain after *Dxz4* inversion, presumably due to a reversal of the mostly unidirectional CTCF binding motifs at *Dxz4* causing new contacts with loci in the opposite direction along the Xi.

Surprisingly, we found an increase in CTCF binding and in chromatin accessibility measured by ATAC-seq, consistent with de-condensation of the Xi in cells with a deletion of the hinge and to a lesser extent in cells with an inversion of *Dxz4.* Consistently, a small increase in Xi relative to Xa volume was observed by microscopy. CTCF binding is normally low on the Xi except at genes that escape XCI^35^, and Minajigi et al. proposed that *Xist* RNA evicts CTCF and cohesin from the Xi^10^. Furthermore, induction of *Xist* expression in male ES cells resulted in the formation of a bipartite structure and loss of chromatin accessibility^9^. However, our results indicate that eviction of CTCF from the Xi in WT and its partial return after *Dxz4* deletion or inversion appear to be independent of *Xist* since there was no change in *Xist* expression or Xi coating. The increase in CTCF binding concomitant with the loss of long-range interactions on the Del-hinge Xi is reminiscent of observations made on neuronal cells after knock-out of the histone methyltransferase SETDB1, which apparently protects the genome of neurons from excessive accumulation of CTCF^36^. One possibility is that loss of long-range contacts and increased chromatin accessibility may facilitate CTCF binding in cells with an alteration of *Dxz4.*

Despite the changes observed in chromatin accessibility in Del-hinge and to a lesser extent in Inv-Dxz4, we observed little reactivation of X-linked genes, even in the presence of a demethylating agent, attesting to the multiple layers of controls that ensure silencing of the X, and suggesting that de-condensation and increased chromatin accessibility are not sufficient to elicit widespread reactivation on the Xi^1,37,38^. Rather, our findings suggest that de-condensation of the Xi due to *Dxz4* deletion may facilitate stochastic dysregulation of a few X-linked genes (Fig. 7b-d). A similar level of clonal variability was reported in a study of mouse ES cells in which expression of genes that escape XCI was reduced after deletion of the hinge and differentiation to neural precursor cells^9^. No changes in the repressive histone mark H3K27me3 enrichment on the Xi were observed in *Dxz4* mutant lines, but we cannot exclude local changes around the locus. Localized loss of H3K27me3 and gain of H3K9me3, as well as a shift in Xi replication timing from early to late, were observed in human cells with a 300kb deletion including *DXZ4,* but there was little change with respect to gene expression^8^. We found no changes in the preferred locations of the Xi near the lamina or the nucleolus in mutant cell lines with a *Dxz4* deletion or inversion, suggesting that additional loci that also bind CTCF and nucleophosmin may facilitate the Xi location^15,30^.

Our findings that the inversion of *Dxz4,* which includes its promoter, causes massive contact redistribution, suggest that the locus itself rather than its lncRNA is critical for the proper orientation of contacts. However, we cannot exclude *Dxz4* transcription as a potentially important factor, possibly by rendering the locus accessible, but this remains to be clarified. The human *DXZ4* and mouse *Dxz4* lncRNAs are expressed from both Xa and Xi alleles^14,15^, but there are differences in the levels and types of transcripts produced from each allele. In mouse, a long sense transcript was detected from the Xa^12^, and we previously reported higher expression in female versus male tissues and cell lines, consistent with Xi expression^15^. In human, the *DXZ4* transcripts include a long sense-transcript and short antisense-transcripts from both alleles, and a long antisense-transcript from the Xi only^39^. The small RNAs produced from the Xa have been hypothesized to help recruit Argonaute proteins to facilitate DNA methylation^39^, which marks the Xa allele and would prevent CTCF binding^12,13,16^.

Analyses of contact distributions show that the greatest level of disruption of the bipartite structure of the Xi after *Dxz4* deletion or inversion is in the vicinity of *Dxz4,* which is in agreement with a study in human where a hinge deletion coincides with the disruption of an interaction compartment (as reflected by principal eigenvectors) in the vicinity of *DXZ4*^8^. A novel finding in our study is that *Dxz4* mainly contacts loci located in the telomeric superdomain and that these long-range contacts are either lost after *Dxz4* deletion or shifted to the centromeric superdomain after *Dxz4* inversion (Fig. 3). CTCF strongly binds at *Dxz4* on the Xi via unidirectional conserved CTCF binding motifs embedded in the tandem repeats of the macrosatellite locus^12,15^. The orientation of CTCF motifs is critical in the formation of chromatin loops by extrusion^11,21,22,25,40^. In the extrusion model, a cohesin ring complex (or other EF) is initially loaded onto two adjacent regions of chromatin in cis and then slides over the chromatin fiber in opposing directions, bringing increasingly remote regions together and resulting in the extrusion of a loop. The processivity of EFs is believed to be curtailed by BEs such as CTCF, apparently in a directional manner, with binding to convergent motifs being far more effective^11,21,22,25,40^. In this scenario, the particularly large bank of Xi-specific CTCF binding at *Dxz4*^12,15^ would present a formidable barrier to EFs, halting chromatin fiber translocation at that end of the loop. At the other end of the loop, EFs would remain relatively free to continue to extrude chromatin, unless they were to encounter another loop or another CTCF binding site preferably in opposite orientation (Fig. 7e). Given the paucity of CTCF binding on the Xi this would result in very large loops. Thus, EFs would tend to stall at *Dxz4,* impeded from moving through to the other side of the locus and effectively insulating the two superdomains from one another. This is consistent with polymer simulations highlighting the role of EFs in imposing insulation over large spatial and genomic distances simply by their translocation processivity being regulated by BEs such as CTCF^21^. Following this reasoning, regions immediately to the rear of *Dxz4* (with directionality dictated by the orientation of the CTCF motifs) would be subject to reduced translocation, due to the EFs being stalled within the bank of CTCF sites. The diminished extrusion would result in fewer long-range contacts and reduced loop formation with chromatin taking on a more unraveled appearance (Fig. 7e). This unraveling ‘behind’ *Dxz4* may also be a result of increased tension in the chromatin fiber due to the locus being pulled closer to *Xist* and *Firre* in the WT and Inv-Dxz4 Xi, respectively (Fig. 3e). The *Dxz4* orientation-dependent shift in contacts is also visible as a telomeric to centromeric switch in the line of strong contacts made by *Dxz4,* a feature that has been termed a “flame”^21,32^, representing strong contacts between a BE and adjacent domain and hypothesized to result from the stalling of extrusion at one end of the loop of DNA in line with the model proposed above (Fig. 1a, 2b, 3b).

Among the new long-range contacts found after *Dxz4* inversion, the most striking is a strong interaction between *Dxz4* and *Firre,* most likely mediated by the banks of CTCF binding sites at both loci^12,15,30^. While the orientation of CTCF motifs is not completely defined at *Firre* due to the repetitive nature of the locus, motif searches indicate that a similar number of CTCF binding sites exist in either orientation. While we only found a small peak of interactions between *Dxz4* and *Firre* in wild-type Patski cells, a previous study reported a stronger peak in human cells^8,11^. The latter could reflect a different configuration in mouse versus human, where a larger *DXZ4* locus contains two sets of CTCF motifs in opposite orientation^14^, which would effectively result in a combination of inverted CTCF binding sites favorable to interactions with loci on both sides including *FIRRE.* Our results suggest that the bank of CTCF binding sites at *Dxz4* still has the ability to curtail the movement of cohesin even when inverted, resulting in a bunching of chromatin fibers and a strong contact domain that does not normally form on the WT Xi (Fig. 7e). Thus, *Dxz4* contacts have little specificity, consistent with the lack of conservation of the *Dxz4* repeat units, except for the CTCF binding motifs and a short sequence (13bp) that could potentially represent a binding site for another BE^12^. A previous study reported that cohesin is excluded from the Xi^10^, but we have reported that at least *Dxz4* and *Firre* strongly bind cohesin on the Xi^15^. Thus, cohesin is not entirely repulsed from the Xi. Further studies are needed to sort out the role of cohesin and CTCF on the Xi. Indeed, loss of CTCF can lead to redistribution of cohesin away from CTCF binding peaks rather than a loss of cohesin^41^.

Our observations are consistent with a model in which *Dxz4* acts as a row of unidirectional obstacles that hook multiple loops of chromatin for condensation with an inherent ability to insulate in either orientation (Fig. 7e). This model also suggests a possible mechanism by which the Xi may fold at the hinge but this remains to be further investigated^6^. The *Dxz4* platform structure may behave in a manner more akin to a ratchet than to velcro, since the locus orientation is important and translocation of chromatin is impeded unidirectionally. In such a scenario cohesin rings would sit along *Dxz4* at sites paired with convergent CTCF sites with chromatin loops threaded through them. Indeed, we and others have reported a strong visible CTCF accumulation within the Xi in interphase nuclei, which co-localizes with DXZ4/Dxz4^42,43^. Although Xi-specific *Dxz4* contacts are represented as occurring simultaneously along the same chromatin fiber in the model and could be interpreted to be stable structures, in reality this is unlikely to be the case. This may well be a dynamic process, with contacts between distant X-linked loci and the *Dxz4* platform being transient albeit relatively frequent. And whether some or all units of the Xi tandem repeat are engaged in loop formation in a single cell is unclear. Single-cell analyses may help better understand cell-to-cell variability in the Xi contact distribution.

## Methods

### Allele-specific CRISPR/Cas9 editing of Patski cells

Patski cells are fibroblasts derived from 18dpc embryonic kidney from a cross between a BL6 female mouse with an *Hprt^BM3^* mutation^44^ and *Mus spretus* male. The cells were selected in HAT media such that the BL6 X chromosome is always inactive as verified in previous studies^28,29^. For allele-specific CRISPR/Cas9 editing sgRNAs designed using CHOPCHOP^45,46^ were selected to include BL6 SNPs at the PAM site if available (Supplementary Fig. S1a and Table S1). Patski cells were transfected using Ultracruz transfection reagents (Santa Cruz). Verifications of the deletions/inversion were done using PCR together with Sanger sequencing to verify specific loss of the BL6 allele with the deletion and to verify junction sequences containing BL6 SNPs (Supplementary Fig. S1b and Table S13). Note that two independent clones (a, b) with a deletion of the hinge and with an inversion of *Dxz4* were derived, while single clones were derived for other deletions or inversion. Del-hinge clone a and Inv-Dxz4 clone a, which have a near-diploid karyotype, were used for all studies, except for gene expression analyses in which we considered both clones a and b.

### Immunostaining and FISH

To verify the presence of a deletion in Del-hinge DNA-FISH was done on metaphase cells using BAC probes (RP23-299L1 for *Dxz4* and RP23-338M16 for *Firre)* as described^15^. For 3D image analysis, DNA-FISH using a whole mouse X chromosome paint probe (XMP X green from MetaSystems) was done in combination with H3K27me3 immunostaining (red) and Hoechst 33342 staining (blue) as described^15^. Images were collected using 21 z-stacks (0.3!m per stack; 10 stacks up and down from the middle plane) using a Nikon TiE inverted wide-field fluorescence microscope. The 3D image analysis was done using Imaris image analysis software for 63, 86, and 51 nuclei from Patski2-4, Del-hinge and Inv-Dxz4, respectively, in which the X-chromosome surface was called based on X-paint probe intensity using the auto-setting threshold. As expected, the condensed Xi marked by H3K27me3 usually showed stronger signals for the X-paint probe compared to the Xa, and also had a stronger Hoechst signal. The Xi volume was calculated and normalized to the Xa volume in the same nucleus (Xi/Xa*15 μm3).

RNA-FISH was done using a 10 kb *Xist* cDNA plasmid (pXho, which contains most of exon 1 of Xist^47^) as described^15^. More than 100 nuclei per genotype were scored for the presence of *Xist* RNA clouds. Immunostaining of paraformaldehyde fixed nuclei using antibodies for H3K27me3 (Upstate/Millipore) and for nucleophosmin (Abcam) was done as described^15^. Nuclei were examined by fluorescence microscopy to score the position of the Xi marked by H3K27me3 with respect to the nuclear periphery and the edge of the nucleolus. For Xi nuclear positioning, a total of ~200 nuclei were scored by at least two different observers.

### Quantitative RT-PCR

RNA extracted from WT, Patski2-4, Del-hinge and Inv-Dxz4 cells by the Qiagen RNeasy kit with on-column DNaseI digestion was reverse-transcribed into first-strand cDNA using SuperScriptII reverse transcriptase (Invitrogen). qPCR was done for two independent cDNA preparations per genotype using gene-specific primers and an ABI7900 qPCR system. The primer sequence information is in Supplementary Table S13.

### In situ DNase Hi-C data analysis

In situ DNase Hi-C was done on intact nuclei from wild-type and deleted/inverted Patski cells using a previously described method^48^. We sequenced the in situ DNase Hi-C libraries using paired-end reads 80bp and 150bp in length (Supplementary Table S2). For each in situ DNase Hi-C library, we mapped each end of the paired-end reads separately to the BL6 genome using the NCBI build v38/mm10 reference genome assembly obtained from the UCSC Genome Browser^49^ and a pseudo-spretus genome using BWA-MEM (v0.7.3) in single-end mode using default parameters^50,51^. A pseudo-spretus genome was assembled by substituting available SNPs (from Sanger Institute, SNP database 2014/10/27 v4) into the BL6 reference genome, as described^35^. We retained only primary reads with MAPQ ≥30 for further analysis. Using heterozygous SNPs between the BL6 genome and the pseudo-*spretus* genome that were validated for our particular Patski cell line, we segregated all high-quality uniquely mapped reads (MAPQ ≥30) using an approach very similar to that described in^7^, with the exception that in order to maximize the number of reads assigned to either the BL6 Xi or the *spretus* Xa, we required that only one end of the read be specifically mapped to one mouse species, while the other end was allowed to be ambiguously mapped. This approach, commonly used to analyze data from hybrid systems, is based on the assumption that intrachromosomal contacts are much more frequent than interchromosomal contacts. Briefly, each end of each read pair was assigned to one of three categories: (1) BL6-SNP reads containing only BL6-specific SNP(s); (2) *spretus*-SNP reads containing only spretus-specific SNP(s); (3) ambiguous reads that did not contain valid SNPs or that contained valid SNPs from both alleles. We refer to both BL6-SNP reads and *spretus-SNP* reads as “allele-specific reads”, and reads that do not contain valid SNPs as “allele-uncertain reads”. Reads were paired with their corresponding mates, and those read pairs with at least one end being allele-specific were retained for subsequent allele-specific analysis. To eliminate the bias due to the PCR duplication step, we removed redundant paired-end reads defined as those pairs where both ends were mapped to identical locations in the same genome assembly. This resulted in a set of valid read pairs representing DNA-DNA interactions (Supplementary Table S2).

We used the resulting valid read pairs to generate allele-specific whole-genome contact maps at 500kb and 40kb resolution. To do so, we partitioned the genome into non-overlapping bins and counted the number of allele-specific contacts (uniquely mapped valid paired-end reads) observed between each pair of bins. The dimension of the resulting allelic contact maps is the total number of bins in the genome, where entry (i, j) contains the contact count between bins i and j. We normalized the allele-specific contact maps using an iterative correction method^52^ to obtain a filtered contact map with near-equal row and column sums. Prior to applying the iterative correction procedure, the contact maps were filtered as follows: bins along the diagonal, super-diagonal (+1 off-diagonal) and sub-diagonal (−1 off-diagonal) (representing entries dominated by self-ligation products) and bins with the lowest 2 % read coverage (representing sparsely populated regions dominated by spurious contacts) had their contact counts set to zero.

For visualization purposes, in order to better compare contact maps across multiple samples, intrachromosomal contact maps were quantile normalized to one another. The Pearson correlation transformation of contact matrices was performed to better visualize the more probable contacts and mitigate for differences in sequencing depth and data sparsity due to allelic-specific read assignment. This transformation was performed as described in^6^, except that the transformation was performed on the ICE-normalized contact matrices rather than the observed-over-expected matrices because we did not wish to normalize out the distance effect.

### Contact decay curves

Plots of the average Hi-C interaction frequencies as a function of genomic distance^6^ were performed using ICE-normalized contact matrices at 500kb resolution either across the length of the chromosomes or within the specified regions of the X chromosome. To account for differences that arise for technical reasons during library preparation, the average autosomal reads per unit of distance from the diagonal for each sample and allele were scale normalized to one another. These autosomal scaling factors were then applied to the X chromosome alleles.

### Coverage score analysis

To count the number of reads spanning each locus along the chromosome across all length scales we made a minor adaptation to the coverage score developed by^31^. Briefly, the coverage score for each bin at a particular resolution was calculated as the average number of interactions within bins spanning this central bin of interest. This score calculation can be visualized as sliding a V-shaped region (with the arms of the V extending all the way to the edges of the contact map) along the diagonal of the contact map and calculating the mean interaction counts (sum of counts/number of bins) within the region. One point of difference compared to Eser and colleagues ^31^ was that instead of normalizing the coverage score for each chromosome to fall within the range [0; 1], we normalized the scores by calculating log2(coverage score/chromosomal mean), as is done in the insulation score calculation (see below). Additionally, we did not restrict our analysis to interactions less than 100kb, but instead considered all interactions spanning each locus (i.e. the arms of the V-shaped region extend all the way to the borders of the contact map). However, when calculating the coverage scores for the unmerged data sets (WT, Del-hinge, Del-Dxz4, Del-Ds-TR, and Inv-5’Ds-TR), we did exclude bins within the first and last 10Mb of the chromosome in order to avoid extreme edge effects that were evident especially in the sparser samples. Furthermore, a five-window, degree two polynomial Savitzky-Golay filter was applied to the resulting vector to smooth the signal, with interpolated values being assigned to edge-case bins. A standardized smoothed coverage score (coverage z-score) or a coverage score rescaled to lie within the [0; 1] interval was used where described in text.

### Insulation score and domain boundary analysis

To determine regions of more localized interaction (domains) and associated boundaries, we used the insulation score method introduced by^33^. Insulation score scripts were downloaded from https://github.com/dekkerlab/cworld-dekker. For 500kb binned DNase Hi-C counts, insulation scores were called using the script matrix2insulation.pl with options --is 3500001 –immean --ss 1000001 --ids 2000001 --nt 0.01 --bmoe 3, while for 40kb data the parameters used were --is 520001 --immean --ss 160001 --ids 320001 --nt 0.01 --bmoe 3. The options have the following definitions:

--is: insulation square size, the size of the insulation square in bp;

--im: insulation mode, the method used to aggregrate signal within insulation square;

--ss: insulation smooth size, the size of insulation vector smoothing vector in bp;

--ids: insulation delta span, the window size around the central bin used to determine insulation change or delta, with the resulting delta vector representing the slope of the insulation score and crossing the horizontal 0 at all peaks and all valleys;

--nt: noise threshold, minimum depth of a valley;

--bmoe: boundary margin of error (specified in number of bins) added to each side of the boundary. TADs were called at 40kb resolution using the script insulation2tads.pl using the default parameters (--mbs 0 -- mts 0, representing minimum boundary strength and minimum TAD strength, respectively). Boundary strength (Fig. 4g) is defined as the difference in the delta vector between the local maximum to the left and local minimum to the right of the boundary bin.

### CTCF ChIP-seq analysis

ChIP-seq was performed on WT and Del-hinge Patski cells using an antibody for CTCF (Upstate/Millipore) and an established protocol^53^. PCR using primers specific for *Dxz4* and *H19* was done to confirm CTCF enrichment (Supplementary Fig. S5a and Table S13). CTCF ChIP and input libraries were sequenced as paired-end reads of 75bp in length (Supplementary Table S4). CTCF and input ChIP-seq paired-end reads were mapped to the BL6 genome using the NCBI build v38/mm10 reference genome assembly obtained from the UCSC Genome Browser^49^ using BWA-MEM (v0.7.3) in paired-end mode using default parameters^50,51^. Primary mapped valid paired reads with MAPQ ≥ 10 were de-duplicated and used for calling CTCF peaks based on biallelic reads using MACS2^54^ (Supplementary Table S4). CTCF peaks were called using all reads (unsegregated) after normalization to their chromatin inputs (Supplementary Table S4). Over 80% of these peaks contained a validated BL6/spretus SNP (Supplementary Fig. S5c), with the vast majority covered by at least 5 reads and considered for allelic analysis (Supplementary Fig. S5d; see Methods). CTCF peaks containing spretus/BL6 SNPs that were covered by a total of at least 5 reads were considered for allelic analysis (Supplementary Fig. S5; Supplementary Table S4). This level of coverage was chosen because the overall allelic proportion (*spretus*/(*spretus*+BL6)) did not change much regardless of whether 5x or 10x coverage thresholds were used (Supplementary Fig. S5f). For each CTCF peak with sufficient SNP coverage, an allelic proportion of SNP read coverage (*spretus*/(*spretus*+BL6)) was calculated. Note that the small difference between autosomal allelic proportion distributions is due to aneuploidy for chromosomes 3 and 4 in some Patski cell clones, and disappears when these chromosomes are excluded from the analysis (Fig. 5a; Supplementary Fig. S5g). Using autosomal distributions of allelic proportion as a guide, peaks with an allelic proportion of greater than 0.7 were designated as spretus/Xa-specific, while those with an allelic proportion of less than 0.3 were deemed to be BL6/Xi-specific. Peaks with an allelic proportion falling within the range [0.3; 0.7] were considered biallelic (Supplementary Fig. S5; Supplementary Table S4). Subtracting one from allelic proportions results in ‘d-scores’ as previously described^9,55^, with values ranging from -0.5 to +0.5. Peaks with positive d-scores are covered by more reads emanating from the *spretus*/Xa allele, while those peaks having negative d-scores show a BL6/Xi bias.

### ATAC-seq analysis

ATAC-seq was done on WT and Del-hinge Patski cells using a published method^56^. ATAC-seq libraries were sequenced in multiple runs as paired-end reads of 75 or 150bp in length (Supplementary Table S5). ATAC-seq read mapping, peak calling, allelic assignment and d-score calculation were performed as described for CTCF peaks, except that ATAC peaks were called without using an input (background) library and all reads were trimmed to 75bp prior to mapping for the sake of consistency (Supplementary Fig. S7; Supplementary Table S5). Fewer ATAC peaks contained SNPs than CTCF peaks (Supplementary Fig. S7a), which could be due to the fact that in both WT and Del-hinge many more, smaller ATAC peaks were called compared to CTCF peaks, possibly a consequence of not having background data to compare against as one does for ChIP-seq with input samples. However, the majority of peaks were covered by at least 5 SNP reads (Supplementary Fig. S7b). The overall allelic proportion (*spretus*/(*spretus*+BL6)) for ATAC peaks did not change much regardless of whether 5x or 10x coverage was required (Supplementary Fig. S7d).

### RNA-seq analysis

RNA-seq was done on WT, Del-hinge (clone a and b), Del-Dxz4, and Del-Ds-TR cells, as well as on Inv-Dxz4 (clone a and b) and the WT subclone Patski2-4 (used to derive Inv-Dxz4) as described^15^. To induce DNA demethylation a 48h treatment using 4!M 5-aza-2’-deoxycytidine (5-aza-2dC) was used, followed by a 24h recovery period, reported to lead to ~2% of cells with reactivation of an X-linked GFP reporter gene^58^. We found that 5-aza-2dC treatment caused ~20% more cell death in WT and Del-hinge cells compared to the mock treatment, similar to that reported for dermal fibroblasts^57^. RNA-seq libraries were sequenced as single-end reads of 75bp in length (Supplementary Table S6). RNA-seq reads were mapped to the UCSC mm10 (NCBI build v38) refSeq transcriptome^49^ as downloaded and packaged in the iGenomes reference sequences and annotation files on July 17, 2015. (https://support.illumina.com/sequencing/sequencing_software/igenome.html). Tophat2 (v 2.0.12) (calling bowtie2 (v2.2.3)) was used to perform single-end mapping allowing 6 mismatches but otherwise default parameters^58,59^. To determine biallelic expression levels, mapped reads were assigned to refSeq genes using HT-seq^60^ and counts were converted into TPMs using custom R scripts.

Genes containing *spretus*/BL6 SNPs that were covered by a total of at least five reads were considered for allelic analysis. For each gene with sufficient SNP coverage, an allelic proportion of SNP read coverage (*spretus*/(*spretus*+BL6)) was calculated. Read counts for each gene were then distributed to each allele based on this SNP-read allelic proportion, allowing us to perform differential expression between samples for each allele. Differential expression analysis was performed using DESeq2^61^. For the WT versus Del-hinge comparison, WT and Del-hinge control samples from the 5-aza-2dC experiment were pooled and treated as biological replicates for the WT and Del-hinge clone a samples.

Genes were deemed to escape XCI in the Patski WT line if their expression levels met the similar criteria to those used by Berletch et al. ^35^ in 2/3 of the WT samples (WT and the two WT 5-aza-2dC untreated replicates) (Supplementary Table S7): (1) the 99% lower confidence limit (alpha = 0.01) of the escape probability was greater than 0.01 based on a binomial distribution parameterized by the expected proportion of reads from the Xi indicating some contribution from the Xi; (2) the diploid gene expression measured by TPM ≥ 1, indicating that the gene was expressed; (3) the Xi-TPM was ≥ 0.1, representing sufficient reads from the Xi, and (4) SNP coverage ≥ 5.

## Data availability

All sequencing data that support the findings of this study have been deposited in the National Centre for Biotechnology Information GEO and are accessible through the GEO SuperSeries GSE59779 as SubSeries GSE107282, GSE107286, GSE107290, GSE107291. All other data and the scripts used for the analyses that support the findings of this study are available from the corresponding authors upon reasonable request.

## Acknowledgements

This study was supported by grants U54DK107979 (JS and WSN) and GM046883 (CMD). We thank C. Lee (University of Washington) for DNA sequencing. We also thank P. Fields and D. Hailey (University of Washington) for their help with 3D imaging and data analysis.

## Supplementary Figure legends

**Supplementary Fig. S1.** Generation and verification of the deletions and inversions of the mouse Xi. **a**. Schematic of the hinge region indicating the position of the guide RNAs used for CRISPR/Cas9 editing. Deletion of the whole hinge (127kb nt75637519-75764753) was obtained in two independent experiments using Ds1 and Dx2 and includes the transcriptional start site of the *Dxz4*-associated lncRNA gene *4933407K13Rik;* Deletion of *Ds-TR* (37kb nt75637501-75674037) was obtained using Ds1 and Ds2 and does not include the promoter region of *Ds-TR;* Inversion (907bp nt75674046-75674952) of 2 of 3 CTCF binding sites located 5’ of *Ds-TR* was obtained using Ds2 and Ds3; Deletion and inversion of *Dxz4* (44kb nt75721096-75764754) were obtained using Dx1 and Dx2 (Supplementary Table S1). **b**. Example of a verification of one of the alterations: PCR amplification using primers F1 and R1 followed by Sanger sequencing verified loss of SNPs from the BL6 allele and PCR amplification using primers F1 and R2 revealed the new junction sequence (see also Supplementary Table S2). **c**. Deletion of the hinge was also verified by fluorescence in situ hybridization (FISH) using a BAC probe for *Dxz4* (red) and a control BAC probe for *Firre* (green). Left, example of a metaphase chromosome preparation with one intact X chromosome with green and red signals (arrows) and a Dxz4-deleted X chromosome with only a green signal (arrow); Right, example of a nucleus with one red and two green signals (arrows).

**Supplementary Fig. S2. a**. Contact maps for the Xa do not differ between WT and Del-hinge, Del-Dxz4, Inv-Dxz4, Del-Ds-TR, and Inv-5’Ds-TR. Contact maps are shown at 500kb resolution. The position of *Dxz4* on the X and schematics of the allele-specific deletions/inversion are shown under the maps (see also Fig. 1a). The color scale reflects the normalized contact counts. **b**. Contact maps for the Xi and Xa representing two pooled sets of Hi-C contacts: one data set representing wild-type and Del-Ds-TR (designated WT*), and the other representing Del-hinge and Del-Dxz4 (designated Del-hinge/Dxz4). Allelic contact maps for each pool are very similar to those obtained from the pooled data (see Fig. 1a for comparison to maps generated for each mutation). Allelic contacts maps for the Inv-Dxz4 Xi and Xa (duplicated in from Fig. 1 and Fig. S2a, respectively) are included to facilitate direct comparisons. Contact maps were generated at 500kb resolution. The location of *Dxz4, Firre* and *Xist* and schematics of the allele-specific deletions/inversions are shown. The color scale reflects normalized contact counts.

**Supplementary Fig. S3.** Allelic principal component (PC) analyses (a-d), differential contact maps (e-h), and 4C analyses (i**). a**. The variance in the PC score profiles for the X chromosomes is explained by the top five allelic principal components for the Xa and Xi for WT*, Del-hinge/Dxz4, and Inv-Dxz4 (see also Fig. 2c, d). **b**. WT*, Del-hinge/Dxz4, and Inv-Dxz4 allelic PC score profiles for an exemplar autosome (chromosome 2) based on distance-corrected, normalized contact maps with counts binned at 500kb resolution (*spretus* top row; B6 bottom row). In each case, the top three allelic PC scores are shown (red, blue and green, respectively). **c**. The variance is explained by the top five allelic principal components averaged across all autosomes chromosomes (*spretus* top row; B6 bottom row) for WT*, Del-hinge/Dxz4, and Inv-Dxz4. **d**. Pairwise spearman correlation values and associated scatterplots between allelic PC1 scores for autosomes concatenated end-to-end for WT*, Del-hinge/Dxz4, and Inv-Dxz4 (see also Fig. 2c-e). **e**. Differential contact map based on untransformed count data at 500kb resolution to highlight differences between Del-hinge/Dxz4 Xi and WT* Xi (loss or gain of contacts in the Del-hinge/Dxz4 versus WT* appear blue or red, respectively). Color scale shows differential normalized contact counts. **f**. As in (d) for differential contact map based on Pearson correlation transformed data. Color scale shows differential Pearson correlation values. **g**. As in (d) to highlight differences between Inv-Dxz4 Xi and WT* Xi. **h**. As in (f) for differential contact map based on Pearson correlation transformed data. **i**. Virtual 4C plots derived from the Hi-C data for various 500kb viewpoints positioned along the Xi for WT* (blue), Del-hinge/Dxz4 (red), and Inv-Dxz4 (black). Y-axis (contact counts) limited to 20% of maximum. The position of the viewpoints is indicated. See text for details and additional data in Fig. 3.

**Supplementary Fig. S4**. Coverage (a-d) and insulation score analyses (e-g). **a**. Standardized coverage Z-score profiles using 500kb resolution Hi-C contact map data along the Xi (top) and Xa (bottom) in WT (blue), Del-hinge (red), Del-Dxz4 (orange), Inv-Dxz4 (black), Del-Ds-TR (green), and Inv-5’Ds-TR cells (indigo). The positions of *Firre, Dxz4,* and *Xist* are indicated (see also Fig. 3d). **b**. Violin plots showing the distribution of standardized coverage Z-scores using 500kb bins for the Xi and Xa in WT (blue), Del-hinge (red), Del-Dxz4 (orange), Inv-Dxz4 (black), Del-Ds-TR (green), and Inv-5’Ds-TR cells (indigo). **c**. Hierarchical clustering based on the Euclidean distance between standardized coverage Z-scores using 500kb bins along the Xi and Xa in WT, Del-hinge, Del-Dxz4, Inv-Dxz4, Del-Ds-TR, and Inv-5’Ds-TR cells. **d**. Hierarchical clustering based on Pearson correlation (using 1 - *r* as the distance measure) for standardized coverage Z-scores using 500kb bins along the Xi and Xa in WT, Del-hinge, Del-Dxz4, Inv-Dxz4, Del-Ds-TR, and Inv-5’Ds-TR cells. **e**. As in (a) for insulation scores. **f**. As in (c) for insulation scores. **g**. Profiles of insulation scores using 40kb bins along the entire length of the Xi (top) and Xa (bottom) in WT* (blue), Del-hinge/Dxz4 (red), and Inv-Dxz4 (black). The positions of *Firre, Dxz4, Zfx, Eda2r,* and *Xist* are indicated (see also Fig. 4).

**Supplementary Fig. S5**. CTCF ChIP-seq SNP coverage and allelic ratios. **a**. PCR amplification of *Dxz4* and of the control imprinted region near the autosomal *H19* gene was done on the input fraction (10% input), the CTCF ChIP fraction (CTCF ChIP) and the no antibody fraction (No Ab) for WT cells (+/+) and Del-hinge cells (Del-hinge/+). A strong decrease in CTCF enrichment at *Dxz4* is seen in Del-hinge cells due to deletion of *Dxz4* on the Xi, which normally binds CTCF. **b**. CTCF binding to *Dxz4* on the Xi is lost in Del-hinge, as shown by ChIP-seq. **c**. Histograms of SNP counts within CTCF peaks along autosomes and the X-chromosomes in WT (blue), Del-hinge (red), and Inv-Dxz4 (grey). **d**. Histograms of the SNP read coverage within SNP-containing CTCF peaks along autosomes and the X-chromosomes in WT (blue), Del-hinge (red), and Inv-Dxz4 (grey). **e**. CTCF peak counts at three different levels of read coverage (0, 5x, 10x) along autosomes and the X-chromosomes in WT (blue), Del-hinge (red), and Inv-Dxz4 (grey). **f**. As in (i) but showing allelic proportions. **g**. Density histograms of the distribution of allelic proportions of CTCF peaks (*spretus/(spretus* +BL6)) along all autosomes and the X-chromosomes for WT (blue), Del-hinge (red) and Inv-Dxz4 (grey). The modes of the X-chromosome distributions are given in parentheses (see also Fig. 5a-c)

**Supplementary Fig. S6**. CTCF and ATAC browser tracks (a, b) and peak location (c, d). **a**. Genome browser tracks of CTCF ChIP-seq peak d-scores ((*spretus*/(*spretus* +BL6) - 0.5) and of peaks assigned as spretus-specific, common, or BL6-specific for WT (blue), Del-hinge (red) and Inv-Dxz4 (black) peaks along the entire X chromosome, and exemplar autosome (chromosome 2), as well as around two imprinted genes (*Peg3, H19*) on chromosome 7. **b**. As in (a) but for ATAC-seq data. **c**. Plots of Xa-specific, common, and Xi-specific CTCF peak density (counts binned within 500kb windows) along the X chromosome for WT (blue), Del-hinge (red), and Inv-Dxz4 (black). To account for differences in the number peaks obtained between samples, the binned counts were scaled by a factor obtained from the between-sample ratios of total diploid autosomal peaks. The scaling factors were 0.7, 0.78 and 1 for WT, Del-hinge, and Inv-Dxz4, respectively. The tables in the top left corner show the total number of peaks under the curves after normalization and those in the top right corner are Spearman correlation matrices across samples (Del = Del-hinge, Inv=Inv-Dxz4). The positions of *Firre, Dxz4,* and *Xist* are indicated. **d**. As in (c) but for ATAC peaks, but the scaling factors were 0.62, 0.63 and 1 for WT, Del-hinge, and Inv-Dxz4, respectively (see also Fig. 7c, d).

**Supplementary Fig. S7**. ATAC-seq SNP coverage and allelic ratios. **a**. Histograms of SNP counts within ATAC peaks along autosomes and the X-chromosomes in WT (blue), Del-hinge (red), and Inv-Dxz4 (grey). **b**. Histograms of the SNP read coverage within SNP-containing ATAC peaks along autosomes and the X-chromosomes in WT (blue), Del-hinge (red), and Inv-Dxz4 (grey). **c**. ATAC peak counts at three different levels of read coverage (0, 5x, 10x) along autosomes and the X-chromosomes in WT (blue) Del-hinge (red), and Inv-Dxz4 (grey). **d**. As in (c) but showing allelic proportions. **e**. Density histograms of the distribution of allelic proportions of ATAC peaks (*spretus*/(*spretus* +BL6)) along all autosomes and the X-chromosomes for WT (blue), Del-hinge (red) and Inv-Dxz4 (grey). The modes of the X-chromosome distributions are given in parentheses (see also Fig. 5d-f).

**Supplementary Fig. S8**. No change of *Xist* expression (a, b) and few changes in gene expression overall upon deletion or inversion of *Dxz4* (c, d). **a**. Expression levels (TPM) of *Xist* from RNA-seq in WT, Patski2-4, Del-hinge clone a and b, and Inv-Dxz4 are similar. **b**. Quantitative RT-PCR confirms similar *Xist* expression levels in WT, Patski2-4, Del-hinge clone a and Inv-Dxz4. No significant change was detected (p-value >0.05 from Student’s t-test). Error bars, s.e.m. *18S* was used for normalization and *ActB* as a control gene. **c**. Plots of expression fold-changes (log2) between Del-hinge versus WT for transcripts on the X chromosome and for subsets of transcripts specifically from the Xi and Xa, based on SNPs, relative to expression levels (mean read counts). The total number of genes examined and the number of genes that show increased expression (Up) or lower expression (Down) in Del-hinge versus WT is indicated. **d**. Plots of expression fold-changes (log2) between Inv-Dxz4 versus wild-type Patski2-4 for transcripts on the X chromosome and for subsets of transcripts specifically from the Xi and Xa, based on SNPs, relative to expression levels (mean read counts). The total number of genes examined and the number of genes that show increased expression (Up) or lower expression (Down) in Inv-Dxz4 versus wild-type is indicated. See additional expression analyses in Supplementary Table S8,S9, S10 and Fig. S9, S10.

**Supplementary Fig. S9**. Gene expression changes in Del-hinge clone a (**a**), Del-hinge clone b (**b**), Del-Dxz4 (**c**), and Del-Ds-TR (**d**) compared to WT cells and Inv-Dxz4 versus wild-type Patski2-4 (**e**). Scatter plots of Xi- and Xa-specific expression between each deleted clone and WT are shown for genes with mean log2(TPM) ≥ -4 diploid (i.e. mean TPM ≥ 0.0625). Dot lines represent 1.5-fold cutoffs. Genes marked by a diamond represent the 16 genes that showed a significant change in Xi-specific expression between WT and Del-hinge clone a (see Fig. 7a). Note that only genes expressed in one condition are labeled.

**Supplementary Fig. S10**. Gene expression changes in WT and Del-hinge clone a after 5-aza-2dC treatment. **a, b**. Plots of expression fold-changes (log2) between 5-aza-2dC -treated cells and controls in WT (a) and Del-hinge clone a (b) for all transcripts on the X chromosome (chrX) and for allele-specific transcripts from the Xi and Xa, relative to expression levels (mean read counts). The total number of genes examined and the number of genes that show increased expression (Up), decreased expression (Down) and the total number of genes with differential expression (DE) in treated versus untreated cells is indicated. **c, d**. Box plots showing the distribution of allelic expression of genes with increased expression (Up) in untreated and 5-aza-2dC treated cells in WT (c) and Del-hinge clone a (d) for autosomal (*A_spretus* or A_BL6) and X-linked (Xa or Xi) genes. Red, *spretus* alleles; blue, BL6 alleles.

**Supplementary Table S1**. List of sgRNAs for CRISP/Cas9 editing. *sgRNAs designed by CHOPCHOP were selected to include BL6 SNPs at the PAM site (red) if available. The *spretus* SNPs are listed as small letters in parenthesis. A pair of sgRNAs were used to edit each target: Ds-1&2 for Del-Ds-TR, Ds-2&3 for Inv-5’ Ds-TR (CTCF peak inversion (chrX:75674066-75674317) at the 5’ of Ds-TR, Dx1&2 for Del-Dxz4 and Inv-Dxz4, and Ds1&Dx2 for Del-hinge. See also Supplementary Fig. S1a.

**Supplementary Table S2**. In situ DNase Hi-C valid read pairs before and after merging. The number of valid read pairs and their length is listed for each Hi-C library together with the calculated total number of reads for each individual cell type, and for the pooled data sets (WT* and Del-hinge/Dxz4). Reads are mapped to mm10 using BWA MEM. The number of BL6-specific and spretus-specific interactions based on one unambiguously mapped end and discarding interactions < 20kb are listed.

**Supplementary Table S3**. Summary of TADs. The estimated number of TADs is shown for the Xi and Xa for each cell line using different bin size.

**Supplementary Table S4**. Summary of CTCF ChIP-seq read counts and peaks. The number of read pairs is listed for the ChIP and the input in Patski WT and Del-hinge, as well as for the ChIP in Patski Inv-Dxz4 and the Patski2-4 clone. The number of diploid CTCF peaks and SNP-containing CTCF peaks including genome-wide, autosomal and X-linked peaks are listed together with the percentage in each category and the ratio between Del-hinge and WT. The number of allelic peaks including covered peaks, Xa- and Xi-assigned peaks and common peaks are listed together with the percentages in each category.

**Supplementary Table S5.** Summary of ATAC-seq read counts and peaks. As described in Supplementary Table S4, but for ATAC-seq.

**Supplementary Table S6**. RNA-seq read counts and alignment statistics. Total RNA-seq reads, mapped reads and their percentage of total are shown for each library.

**Supplementary Table S7**. Genes (mm10) called escape in 2/3 Patski WT lines. The genes listed were classified as genes that escape XCI in 3/3 WT lines based on criteria described in the text, except for *Ftx* classified as escape in 2/3 lines. For each line, the ratio between the number of reads from the Xi versus the total number of reads assigned either to the Xi or Xa is listed, together with the number of reads containing SNPs specific for the Xi or Xa, the total TPM, the Xi- and Xa- specific TPM, and the lower and upper confidence limits of escape probability (see methods).

**Supplementary Table S8**. Differential Xi-specific expression changes between WT and Del-hinge clone a. DESeq2 differential expression results between Del-hinge clone a and WT for each X-linked gene. Columns give the mean expression, log2 fold change in expression, standard error in log2 fold change, DE test statistic, *p*-value of test, and p-value adjusted to account for multiple testing. Highlighted genes show a significant differential expression (|log fold change| > 0.5 and adjusted *p*-value < 0.05). The additional columns give the Xi- and Xa-specific TPM values for each of the replicates used in the differential analysis.

**Supplementary Table S9**. Differential autosomal gene expression changes between WT and Del-hinge clone a. As described in Supplementary Table S8, but only for 2689 significantly differentially expressed autosomal genes based on biallelic expression. TPM values not included.

**Supplementary Table S10**. Differential Xi-specific expression changes between wild-type Patski2-4 and Inv-Dxz4. As described in Supplementary Table S8, but for Inv-Dxz4 and Patski2-4.

**Supplementary Table S11**. Differential Xi-specific expression changes between 5-aza-2dC untreated and treated WT. As described in Supplementary Table S8, but for 5-aza-2dC treated and untreated WT.

**Supplementary Table S12**. Differential Xi-specific expression changes between 5-aza-2dC untreated and treated Del-hinge clone a. As described in Supplementary Table S8, but for 5-aza-2dC treated and untreated Del-hinge clone a.

**Supplementary Table S13**. List of primers. Five different combinations of primer pairs were used for each pair of CRISPR/Cas9 cutting sites to verify deletion or inversion of loci. For example, for *Dxz4* cut1 and 2, Dx_F1/R1, Dx_F2/R2, Dx_F1/R2 were used to verify deletion, and Dx_F1/F2 to verify inversion (see also Fig. S1a, b). Primers to verify the efficiency of CTCF ChIP are listed for *Dxz4* and *H19.*

